# Aerobic metabolism in *Vibrio cholerae* is required for population expansion during infection

**DOI:** 10.1101/2020.06.16.155382

**Authors:** Andrew J. Van Alst, Victor J. DiRita

**Author notes:** Corresponding author Mailing Address: Department of Microbiology & Molecular Genetics, Michigan State University, 2215 Biomedical & Physical Sciences Building, 567 Wilson Road, East Lansing, 48824.

## Abstract

*Vibrio cholerae* is a bacterial pathogen that replicates to high cell density in the small intestine of human hosts leading to the diarrheal disease cholera. During infection, *V. cholerae* senses and responds to environmental signals that govern cellular responses. Spatial localization of *V. cholerae* within the intestine affects nutrient availability and therefore the metabolic pathways required for the replicative success of the pathogen. Metabolic processes used by *V. cholerae* to reach such high cell densities are not fully known. Here we seek to better define the metabolic traits that contribute to high levels of *V. cholerae* during infection by investigating mutant strains in key carbohydrate metabolism pathways. By disrupting the pyruvate dehydrogenase (PDH) complex and pyruvate formate-lyase (PFL), we could differentiate aerobic and anaerobic metabolic pathway involvement in *V. cholerae* proliferation. We demonstrate that oxidative metabolism is a key contributor to the replicative success of *V. cholerae in vivo* using an infant mouse model where PDH mutants were attenuated 100-fold relative to wild type for colonization. Additionally, metabolism of host substrates such as mucin were determined to support *V. cholerae* growth *in vitro* as a sole carbon source primarily in aerobic growth conditions. Mucin likely contributes to population expansion during human infection as it is a ubiquitous source of carbohydrates. These data highlight the importance of oxidative metabolism in the intestinal environment and warrants further investigation of how oxygen and other host substrates shape the intestinal landscape that ultimately influences bacterial disease. We conclude from our results that oxidative metabolism of host substrates such as mucin is a key driver of *V. cholerae* growth and proliferation during infection, leading to the substantial bacterial burden exhibited in cholera patients.

**Importance:** *Vibrio cholerae* remains a challenge in the developing world and incidence of the disease it causes, cholera, is anticipated to increase with rising global temperatures and with emergent, highly infectious strains. At present, the underlying metabolic processes that support *V. cholerae* growth during infection are less well understood than specific virulence traits such as production of a toxin or pilus. In this study we determined that oxidative metabolism of host substrates such as mucin contribute significantly to *V. cholerae* population expansion *in vivo*. Identifying metabolic pathways critical for growth can provide avenues for controlling *V. cholerae* infection and the knowledge may be translatable to other pathogens of the gastrointestinal tract.

## Introduction

*Vibrio cholerae* causes the diarrheagenic disease cholera in humans and is particularly problematic in regions of the world with poor water sanitation. Ingesting contaminated water sources containing sufficiently high numbers of *V. cholerae* bacterial cells leads to infection characterized by excessive fluid loss and a substantial bacterial burden of *V. cholerae* during the acute phase of disease. In the human gastrointestinal tract, *V. cholerae* can proliferate to numbers as high as 10^6^ to 10^8^ cells per gram of stool (1). Here we seek to understand the metabolic requirements for *V. cholerae* that support such substantial population expansion within the gut.

The mucus lining of the gastrointestinal tract, which typically serves as a barrier to infection, is saturated with a variety of carbohydrates. Mucin is a glycoprotein and the primary macromolecule of mucus. Mucin consists of a protein backbone decorated with *O*-linked glycan chains containing sugars such as *N*-acetylgalactosamine, *N*-acetylglucosamine, galactose, fucose, and sialic acid (2). Commensal mucin-degrading bacteria such as *Bacteroides* spp. And *Akkermansia mucinophila* contain numerous mucinolytic enzymes capable of releasing these sugars from the mucin glycan chain to support growth (2, 3). Mucin degradation is also a feature of bacterial pathogens such as *Shigella flexneri, Helicobacter pylori*, and enterohemorrhagic *Escherichia coli* (4–7). *V. cholerae* contains a number of mucolytic glycosyl hydrolases that are predicted to release glycans from mucin polysaccharides (8–10). Indeed, previous studies have linked mucus carbohydrate metabolism with infection as *V. cholerae* mutants defective for *N*-acetylglucosamine and sialic acid metabolism were attenuated for colonization in the infant mouse (9–11). The mechanism of acquisition of these mucin carbohydrates may be through both phosphoenolpyruvate-phosphostransferase dependent and independent systems to support growth *in vivo* (12). Although mucin can serve as a substrate for growth, chemical reduction of intestinal mucus during infection leads to increased numbers of *V. cholerae*, indicating that mucus also contributes to intestinal protection and clearance of the bacteria (13).

*V. cholerae* harbors the complete enzymatic pathways for the Embden-Meyerhof-Parnas pathway (EMP/glycolysis), the Entner-Doudoroff pathway (ED), and the pentose phosphate pathway (PP) (14, 15). The EMP and ED pathways predominantly generate the energy necessary for *V. cholerae* growth and proliferation. Additionally, previous work has shown that these pathways promote virulence factor production, although the direct cause for this effect is unknown (15, 16). In contrast, the PP pathway does not appear to play a significant role in the growth or colonization of *V. cholerae* during infection (17). These pathways culminate with the formation of pyruvate which can then be used by the bacterium to fuel either aerobic or anaerobic metabolism to generate energy for the cell.

To expand our understanding of carbohydrate metabolism and its impact on *V. cholerae in vivo* fitness, we targeted the pyruvate dehydrogenase (PDH) complex and pyruvate formate-lyase (PFL) that both function to convert pyruvate to acetyl-CoA (18, 19). Examination of PDH and PFL mutants enables us to assess the contribution of aerobic and anaerobic metabolism to the expansion of *V. cholerae* during infection. The conversion of pyruvate to acetyl-CoA precedes the TCA cycle, as the first step in the cycle requires acetyl-CoA to generate citrate. In previous work, *V. cholerae* mutants defective in the TCA cycle expressed increased levels of *toxT*, which encodes the major virulence gene activator in *V. cholerae*. This finding suggested a link between acetyl-CoA and virulence expression (20). However, these TCA cycle mutants were not tested *in vivo* and have been investigated only in classical biotype strains, not in strains of the El Tor biotype. Classical *V. cholerae* predominated among epidemic isolates prior to 1961, when it was supplanted by the El Tor biotype (21). The biotypes are differentiated by numerous physiological attributes that contributed to displacement of the classical by the El Tor (22–25). Some of these are encoded on genomic islands unique to the El Tor biotype that contribute to phage resistance or acquisition of substrates (26, 27).

Here we assess pathways of carbohydrate metabolism on growth, virulence factor production, and colonization of *V. cholerae* El Tor strain C6706. By targeting the PDH complex and PFL, we are able to draw conclusions about the aerobic and anaerobic metabolic processes that facilitate population expansion of *V. cholerae* during infection. Our results provide evidence supporting the importance of a functional PDH complex during infection, with significantly less reliance on PFL function. This indicates that oxygen metabolism primarily drives the growth and proliferation required to amass the high bacterial cell density observed during the disease cholera. The defects in colonization observed with strains lacking a functional PDH are attributable primarily to metabolic deficiencies as virulence factor production was unaffected by mutation in these metabolic pathways. Given what is known in regard to oxygen availability within the intestinal environment, being highest in the intestinal crypts and decreasing to near hypoxia in the lumen, we can deduce the biogeographical localization of replicative *V. cholerae* (28). Our work suggests that replication within intestinal crypt spaces, observed in findings of others (13), is enabled due to the higher oxygenation of this site compared to the lumen. Furthermore, by using the physiologically relevant growth substrate mucin, we could closely reflect, and assess, the growth substrates typically encountered by *V. cholerae* during infection. The results of this study further our understanding of central metabolism and its contribution to *V. cholerae* infectivity and *in vivo* growth.

## Material and Methods

### Transposon mutagenesis library screen

We used a non-redundant transposon mutant library collection constructed in the El Tor C6706 background (29). Using a 96-well plate replicator, the library was replica plated onto large LB + kanamycin (0.05mg/mL) agar plates and incubated overnight at 37°C. Subsequently, this LB grown library was replica plated onto minimal MCLMAN media (30) plates supplemented with 0.5% Type III porcine gastric mucin (Sigma). Transposon insertion mutants that were qualitatively defective for growth compared to neighboring transposon insertion mutants were marked as deficient for mucin utilization for further investigation. The complete list of identified mutants is in Supplementary Table 1.

### Porcine small intestinal mucus collection and mucin purification

Fresh porcine small intestinal segments were harvested from healthy adult pigs from the Michigan State University Meat Lab. Mucus was scraped from the intestinal segments and purified similarly as previously described (31). Briefly, crude mucus was solubilized and resuspended in Extraction guanidine hydrochloride (Extraction GuHCl) (6M guanidine hydrochloride, 5mM EDTA, 0.01M NaH_2_PO_4_, pH 6.5) and homogenized using a dounce homogenizer. The crude mucus was then rocked overnight at 4°C followed by centrifugation at 14000rpm, 10°C, for 45min. Supernatant was removed, and samples washed again with Extraction GuHCl. Samples were washed and centrifuged a total of five times or until the supernatant appeared clear for two consecutive washes. Mucin was then solubilized using 20mL Reduction guanidine hydrochloride (Reduction GuHCl) (6M guanidine hydrochloride, 0.1M Tris, 0.5mM EDTA, pH 8.0) with the addition of 25mM dithiothreitol (DTT) as powder just before use and rocked for 5 hours at 37°C. 75mM iodoacetamide was added after incubation as powder and samples rotated in the dark overnight. Samples were then centrifuged at 4000rpm for 45min at 4°C. The supernatant was added to dialysis tubing and dialyzed in ddH_2_O for a total of six changes. The samples were flash frozen using liquid nitrogen and lyophilized for purified mucin powder.

### Bacterial strains and growth conditions

*Vibrio cholerae* and *Escherichia coli* strains used in this study are listed in Supplementary Table 2. Unless otherwise specified, *V. cholerae* and *E. coli* strains were grown aerobically at 37°C on LB agar plates or shaking at 210rpm in LB broth. Where indicated, antibiotics were routinely added to the media at concentrations: 0.1mg /mL streptomycin, 0.1mg/mL ampicillin, kanamycin 0.05mg/mL. LB media was prepared according to a previously reported recipe (32), however, solid media was made with a 1.5% w/v concentration of agar.

### Primers

Primers used in this study are listed in Supplementary Table 3.

### Plasmid construction

Plasmid construct inserts were generated by PCR using Phusion high-fidelity polymerase (Thermo Scientific). Vector backbones were generated by plasmid purification using Qiagen Mini Prep Kit and subsequent restriction digest.

A modified pKAS32 suicide vector was constructed to generate *ΔaceE, ΔaceF*, and *ΔpflA* strains (33). Primer sets were used to amplify 1000bp homologous regions upstream and downstream of the target gene. The pKAS32 vector was restriction digested using SacI and XbaI at 37°C for 1 hour followed by an additional 30 minutes at 37°C with alkaline phosphatase from calf intestine (CIP) (New England Biolabs). Vector backbone and upstream and downstream segments were joined using Gibson assembly (New England Biolabs) and subsequently electroporated into electrocompetent S17 λpir *E. coli* and recovered on LB + 0.1mg/mL ampicillin agar plates.

Construction of complementation plasmids can be found in the Supplemental Methods.

### *Vibrio cholerae* mutant construction

Wild type *V. cholerae* and *E. coli* strains were mated on LB agar plates at 37°C overnight. The mating was then plated on LB + 0.1mg/mL ampicillin + 25 U/mL polymyxin B. Colonies were then subjected to streptomycin counter-selection as described previously using LB + 2.5mg/mL streptomycin counter-selection plates (33). Colonies were screened for the deletion using primer sets upstream and downstream of the pKAS32 homology regions.

Generation of complementation strains can be found in the Supplemental Methods.

### Growth curves

M9 + 0.5% purified porcine small intestinal mucin (PSIM) was made by combing in a 1:1 mixture 2X M9 minimal media and 2X (1%) PSIM prepared as a final 20mL volume which was then autoclaved for 20 minutes at 121°C.

Strains were initially grown on LB + 0.1mg/mL streptomycin plates overnight at 37°C and a single colony isolate used to start a fresh LB + 0.1mg/mL streptomycin broth culture grown overnight 210rpm at 37°C. Overnight cultures were washed twice in PBS and resuspended to an optical density 1.0 OD_600_.

#### Aerobic growth curves

For LB growth curves, a 1:1000 dilution of a 1.0 OD_600_ culture was used to inoculate prepared media (either 2mLs or 50mLs, depending on the experiment) which was grown at 210rpm at 37°C. For M9 + 0.5% PSIM, 2mL of media was added to a 15mL round bottom tube and inoculated 1:250 with 1.0 OD_600_ culture and grown 210rpm at 37°C. At each timepoint, 100µl was removed for dilution series plating.

#### Anaerobic growth curves

Anaerobiosis was achieved using a Coy Anaerobic Chamber. For LB growth curves, a 1:1000 dilution of a 1.0 OD_600_ culture was used to inoculate prepared media which was grown statically at 37°C. WT, *ΔaceE*, and *ΔaceF* growth curves were performed in 50mL of LB media in a 125mL flask, whereas later WT and *ΔpflA* LB growth curves were performed in 2mL of LB media in a 15mL round bottom tube. For M9 + 0.5% PSIM, 2mL media was added to a 15mL round bottom tube and inoculated 1:250 with 1.0 OD_600_ culture and grown statically at 37°C. At each timepoint, flask and tubes were swirled or vortexed and 100µl was removed for dilution series plating. For growth curves including 50mM fumarate, sodium fumarate (Sigma) reagent was used.

#### Complementation growth curves

Methods for complementation growth curves can be found in the Supplemental Methods.

### AKI virulence-inducing conditions

#### Standard AKI Conditions

Wild type, *ΔaceE, ΔaceF*, and *ΔtoxT* strains were grown statically in 50mL pre-warmed AKI media in 50mL conical tubes for 4 hours at 37°C followed by a transfer to 125mL flasks, and shaking 210rpm at 37°C (34). 1mL of media was removed at each timepoint and centrifuged 14000rpm for 1 minute. Supernatant was separated from the pellet and stored at −80°C for cholera toxin quantification. The bacterial pellet was resuspended in 1mL TRIzol (Ambion Life Technologies) and stored at −80°C for RNA isolation.

#### Anaerobic AKI conditions

Wild type, *ΔaceE, ΔaceF*, and *ΔtoxT* strains were grown in 50mL pre-warmed oxygen-depleted AKI media in 50mL conical tubes statically in anaerobic conditions using a Coy Anaerobic Chamber (35). 1mL of media was removed at each timepoint and centrifuged 14000rpm for 1 minute. Supernatant was separated from the pellet and stored at −80°C for cholera toxin quantification.

### Cholera toxin quantification by ELISA

Cholera toxin in *V. cholerae* supernatants from standard and anaerobic AKI conditions was quantified by GM1 ELISA as previously described (36, 37). GM1 coated microtiter plates were incubated with a 1:20 dilution of culture supernatant and detected using primary anti-cholera toxin and secondary HRP-conjugated goat anti-rabbit IgG (Invitrogen). 1-Step Ultra TMB-ELISA (Thermo Scientific) reagent was added and stabilized using 2M sulfuric acid. Colorimetric measurements were read at 450nm and the toxin concentration was determined by comparison to a standard curve using purified cholera toxin.

### RNA isolation and Real Time Quantitative PCR (RT-qPCR)

RNA was harvested from AKI toxin-inducing culture pellets preserved in 1mL Trizol using an RNEasy kit (Qiagen) using both on-column DNase digestion (Qiagen) followed by Turbo DNase digestion (Invitrogen). RNA concentration and quality were measured by Nanodrop and visualized on a 2% agarose gel.

Complementary cDNA was generated from RNA using Superscript III Reverse Transcriptase (Thermo Scientific). RT-qPCR reactions were carried out using SYBR Green Master Mix (Applied Biosystems) with 5ng cDNA. Primers used to detect *recA, toxT, ctxA*, and *tcpA* transcripts are listed in Supplementary Table 3. ΔΔCt values were calculated using *recA* as the gene of reference (38).

### Infant mouse colonization assays

All animal experiments in this study were approved by the Institutional Animal Care and Use Committee at Michigan State University.

Infant mice were infected as described previously (39). Three-to five-day old CD-1 mice (Charles River, Wilmington, MA) were oro-gastrically inoculated with approximately 10^6^ bacterial cells two hours after separation from the dam and maintained at 30°C. Mice were euthanized approximately 20 hours after inoculation. Mouse intestinal segments were weighed and homogenized in 3mL PBS. Intestinal homogenates were serially diluted and plated on LB + 0.1mg/mL streptomycin for mono-associated infections and LB + 0.1mg/mL streptomycin + 0.08mg/mL 5-bromo-4-chloro-3-indolyl-β-D-galactopyranoside (X-Gal) for competition infections. Competition infections consisted of a 1:1 mix of target strains with a *ΔlacZ* strain for differentiation by blue-white screening.

For PDH mono-associated and competition infections, the entire intestinal tract was homogenized for bacterial enumeration. In intestinal segment measurements, approximately 1cm of intestine from each section (proximal, medial, and distal) was homogenized for bacterial enumeration between segments. For PFL mono-associated and competition infections, the intestinal tract was divided into small intestine and large intestine plus cecum. The divided intestinal portions were then homogenized for bacterial enumeration.

### Statistical methods

For determining the relationship between cholera toxin output versus optical density among WT, *ΔaceE*, and *ΔaceF* strains, a simple linear regression and subsequent slope and intercept analysis was performed using GraphPad PRISM software, which follows a method equivalent to an Analysis of Covariance (ANCOVA).

For *in vivo* experiments, CFU/g intestine and competitive index scores were log_10_-transformed and tested for normality using a Shapiro-Wilks test. Normally distributed data were then analyzed using either parametric Student’s t-test or an Analysis of Variance (ANOVA) with post-hoc Tukey’s test to test for significance.

For *in vivo* intestinal segment data where bacterial loads were below the limit of detection, a non-parametric Kruskal-Wallis one-way analysis of variance was used with post-hoc Dunn’s test to test for significance.

## Results

### Transposon mutagenesis screen identified the pyruvate dehydrogenase complex as important for growth on mucin

We hypothesized that intestinal mucin would serve as a growth substrate for *V. cholerae* during colonization. In a pilot experiment, *V. cholerae* was observed to exhibit enhanced growth in minimal media supplemented with mucin (Supplementary Figure S1). We then performed a transposon mutant library screen of *V. cholerae* El Tor strain C6706 on minimal media supplemented with 0.5% mucin (Type III Sigma). Genes encoding two of the three components of the pyruvate dehydrogenase (PDH) complex were identified in our screen, *aceE* (VC2414) and *aceF* (VC2413) (Supplementary Table 2). The third component of the PDH complex, *lpdA* (VC2414), was also defective for growth in our screen, however, growth of this transposon mutant was also severely attenuated for growth on LB as this enzyme also functions in the alpha-ketoglutarate dehydrogenase (AKGDH) and glycine cleavage multi-enzyme (GCV) systems (40). Because of its pleiotropic growth defect, we did not further investigate an *lpdA* mutant.

### The pyruvate dehydrogenase complex supports aerobic growth on mucin

As a glycoprotein, mucin is coated in glycans, contributing to the protective function of the mucus barrier (41). To study mucin from a physiologically relevant site of infection, we purified it from the small intestine of healthy adult pigs using a guanidine hydrochloride extraction procedure. The purified mucin was then analyzed by High-Performance Anion-Exchange Chromatography coupled with Pulsed Amperometric Detection (HPAEC-PAD) (GlycoAnalytics) (42, 43). Mucin obtained by this method showed high relative presence of mucin carbohydrates galactosamine, glucosamine, galactose, fucose, and sialic acids Neu5Ac and Neu5Gc compared to non-mucin monosaccharides glucose and mannose (Supplementary Table 4).

When grown in M9 minimal salts media supplemented with 0.5% purified small intestinal mucin (PSIM), isogenic PDH mutants *ΔaceE* and *ΔaceF* were defective for growth in aerobically grown cultures compared to wild type (Figure 1A). This phenotype was complemented for both *ΔaceE* and *ΔaceF* using the isopropyl β-D-1-thiogalactopyranoside (IPTG) inducible pMMB66EH vector in M9 0.5% glucose media (Supplementary Figure S2). To verify the PDH complex does not contribute to anaerobic proliferation on mucin, we measured growth in anaerobic conditions. When grown anaerobically, PDH mutants grew comparable to wild type (Figure 1B). To determine if addition of an alternative electron acceptor would elicit a growth difference between wild type and PDH mutants, 50mM fumarate was added to the growth media, which enhances *V. cholerae* growth via anaerobic respiration (44). No growth disparity was observed between wild type and PDH mutants with the addition of 50mM fumarate (Figure 1C). The observed phenotypes indicate that the PDH complex is not required for anaerobic growth. Energy generation under anaerobic conditions is likely due to an active pyruvate formate-lyase converting pyruvate to formate and acetyl-CoA for mixed acid fermentation (45) and acetolactate synthase that converts pyruvate to (*S*)-2-acetolactate in the first step of 2,3-Butanediol fermentation (25). The growth disparity observed in the minimal mucin media under aerobic conditions was less pronounced in LB media, which contains less than 100µM of collective sugars and primarily supports growth through amino acid catabolism (Supplemental Figure S3) (46).

**Figure 1.**
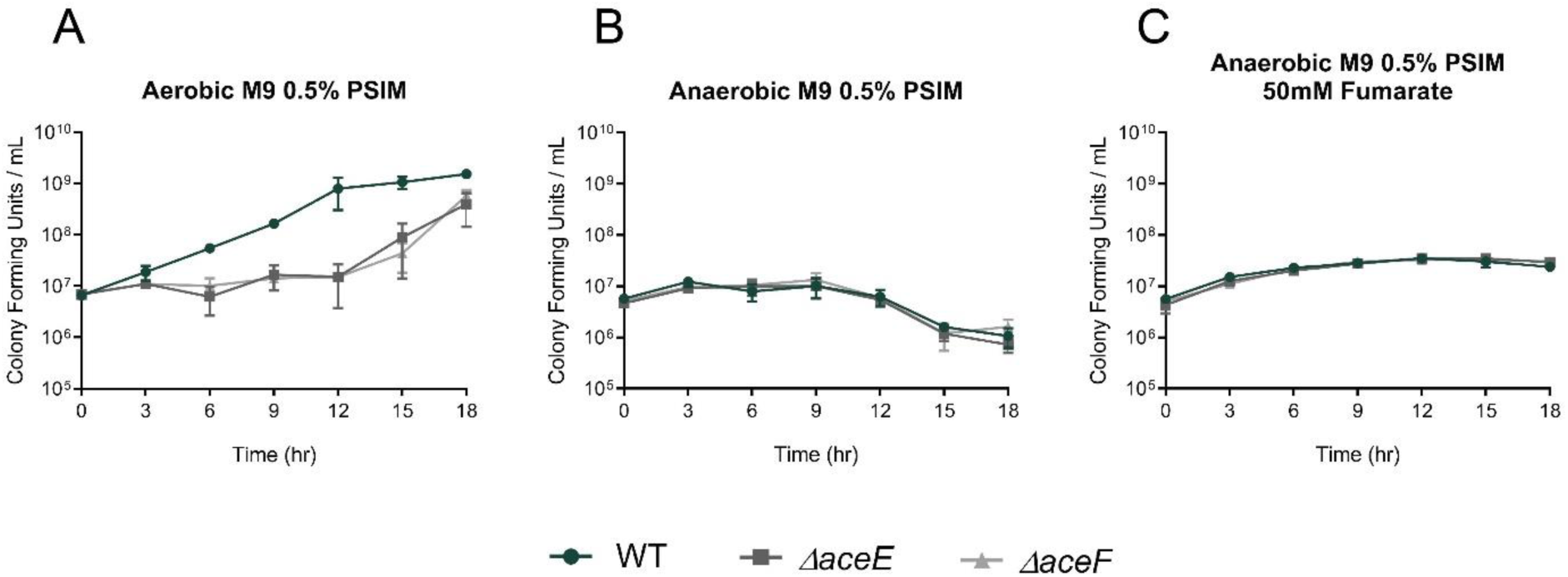
Growth curves of WT (green circle) *ΔaceE* (dark grey square) and *ΔaceF* (grey triangle) in M9 minimal media supplemented with 0.5% purified porcine small intestinal mucin (PSIM) grown (A) aerobically, (B) anaerobically, or (C) anaerobically supplemented with 50mM fumarate. Data represent the average and SEM for three independent biological replicates.

### Pyruvate formate-lyase supports anaerobic growth on mucin

As pyruvate formate-lyase (PFL) also converts pyruvate to acetyl-CoA, we sought to investigate the role of PFL in the catabolism of mucin. To accomplish this, an isogenic mutant strain deleted for *pflA* (VC1869) was tested for *in vitro* growth on PSIM. When grown in M9 minimal salts media supplemented with 0.5% PSIM, *ΔpflA* grew comparable to wild type in aerobic growth conditions (Figure 2A) and poorly compared to wild type (which did not thrive itself) when cultured anaerobically, both in the absence and presence of 50mM fumarate (Figure 2B-C). This growth defect was complemented for *pflA* using an IPTG inducible pMMB66EH vector in M9 minimal salts 0.5% glucose 50mM fumarate media (Supplementary Figure S4). The disparity in growth between wild type and *ΔpflA* strains under anaerobic growth conditions indicates that PFL can indeed generate energy from mucin during anaerobic growth. As PFL is expected to function primarily in the metabolism of carbohydrates it was not surprising to see growth comparable to wild type in LB media, both aerobically and anaerobically, but it was intriguing to find *ΔpflA* remained viable longer than wild type in LB media without the addition of 50mM fumarate (Supplementary Figure S5).

**Figure 2.**
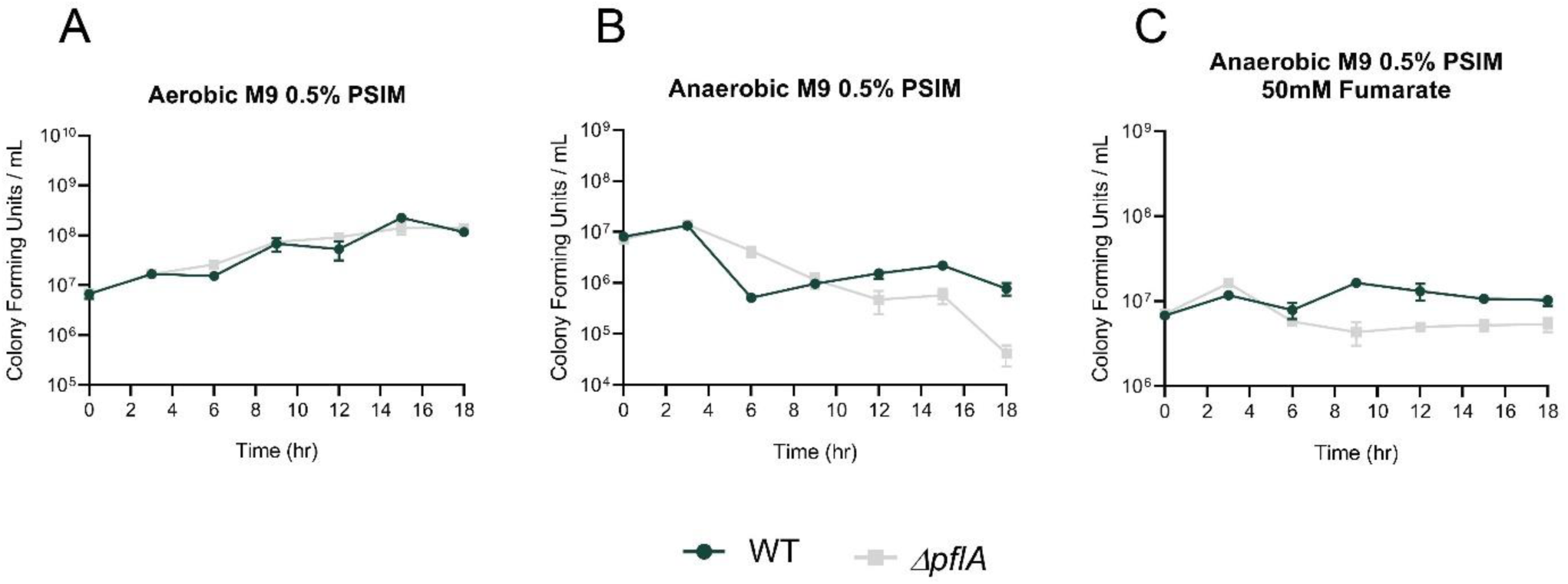
Growth curves of WT (green circle) and *ΔpflA* (light grey square) in M9 minimal media supplemented with 0.5% purified porcine small intestinal mucin grown (A) aerobically, (B) anaerobically, or (C) anaerobically supplemented with 50mM fumarate. Data represent the average and SEM for three independent biological replicates.

### Cholera toxin production in PDH mutants is equivalent to wild type in both standard and anaerobic toxin-inducing conditions

Cholera toxin is the primary virulence determinant of *V. cholerae*. To determine whether the PDH complex influences production of cholera toxin, wild type and PDH mutant strains were grown in AKI conditions to induce virulence factor production (34). Previous findings with strains of the classical biotype demonstrate that disruption of the TCA cycle increases *toxT* expression and suggests a link between acetyl-CoA and virulence expression (20). As the PDH complex is the primary enzyme responsible for the production of acetyl-CoA under aerobic growth conditions, we anticipated mutants lacking it would produce cholera toxin levels below that of wild type. However, we observed no significant difference in cholera toxin produced in the WT and PDH mutant strains in either standard or anaerobic AKI conditions. For standard AKI conditions, cholera toxin levels were measured as a function of optical density with *ΔtoxT* (VC0838) included as a negative control as ToxT stimulates cholera toxin production (Figure 3A) (47). Overall, wild type produced more total cholera toxin as it reached a higher final optical density than PDH mutants, yet when determining individual cellular capacity for cholera toxin production, at similar optical densities, cholera toxin output in the PDH mutants was comparable to wild type. Cholera toxin production levels at each timepoint 4h, 5h, 6h, 7h, 8h, and 24h are provided in Supplemental Figure S6. Cholera toxin levels were also measured after 8h and 20h of growth in anaerobic AKI conditions and were again similar between wild type and PDH mutant strains (Figure 3B and Supplemental Figure S7). These findings indicate that the PDH complex does not affect cholera toxin production in El Tor C6706 *V. cholerae*.

**Figure 3.**
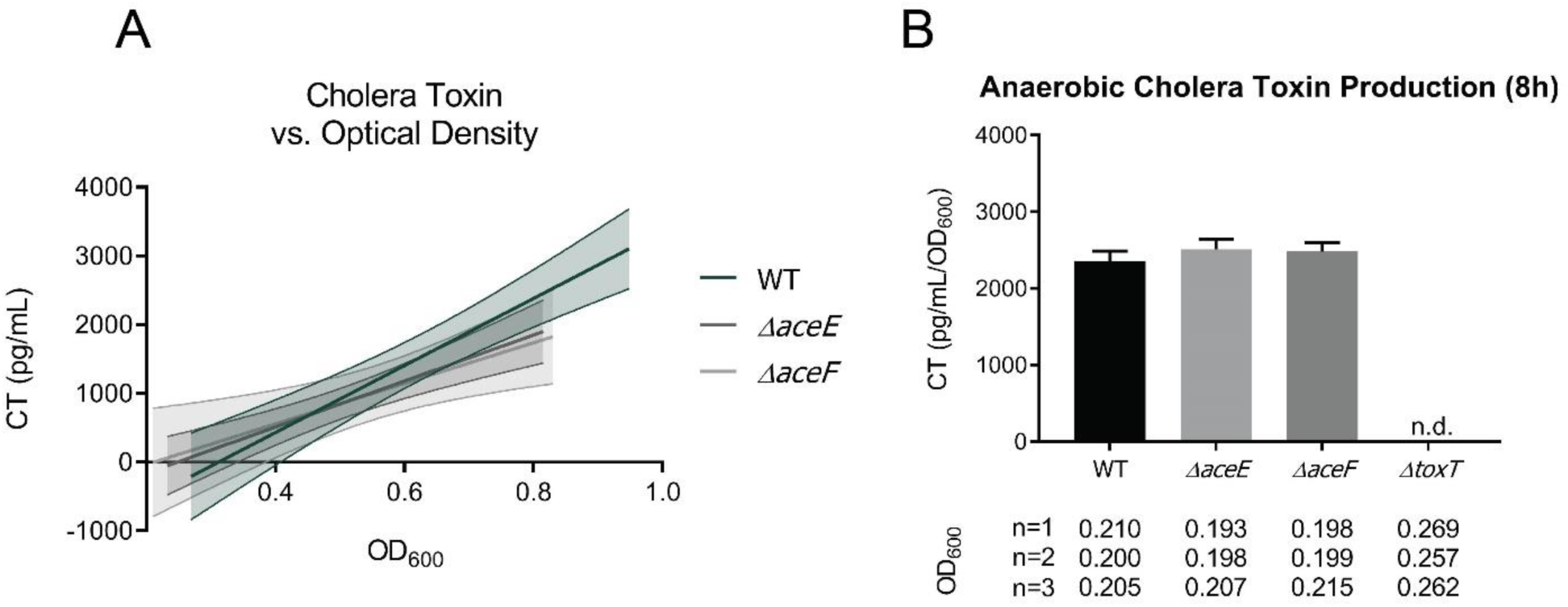
Cholera toxin (CT) production for WT, *ΔaceE, ΔaceF*, and *ΔtoxT* strains. (A) CT output as a function of optical density (OD_600_) under standard AKI toxin-inducing conditions. Data points were collected from three biological replicates and a line of best fit with 95% confidence intervals was plotted. WT OD’s higher than 0.9 were excluded to better superimpose with *ΔaceE* and *ΔaceF* OD values. A simple linear regression found no significant differences between WT, *ΔaceE*, and *ΔaceF* CT production. *ΔtoxT* control was not plotted because no toxin was detected. Statistical analysis was performed using GraphPad PRISM (B) CT values relative to optical density (pg/mL/OD_600_) are reported under anaerobic AKI toxin-inducing conditions at the 8h timepoint. The optical densities for the biological replicates are displayed below the corresponding strain on the x-axis in each graph. Data represent the average and SEM for three biological replicates.

### Functional PDH activity is not required for expression of *toxT, ctxA* and *tcpA*

The relative expression of the master virulence regulator *toxT* and primary virulence factors *ctxA* and *tcpA* was determined by RT-qPCR. The 4h and 5h timepoints of *in vitro* standard AKI conditions were selected to compare relative expression profiles as *toxT* expression has been observed to be high at these timepoints (48). PDH mutant strains at 4h exhibited somewhat elevated levels of *toxT* and *tcpA* and lower *ctxA* expression compared to wild type, with no transcripts detected in the *ΔtoxT* control as expected (Figure 4A). At 5h, PDH mutant strains exhibited wild type *toxT* and *ctxA* expression and a two-fold reduction in *tcpA* expression (Figure 4B). Although there is variability in the expression of these virulence genes at the given timepoints, we conclude that the PDH complex is not required for virulence gene expression.

**Figure 4.**
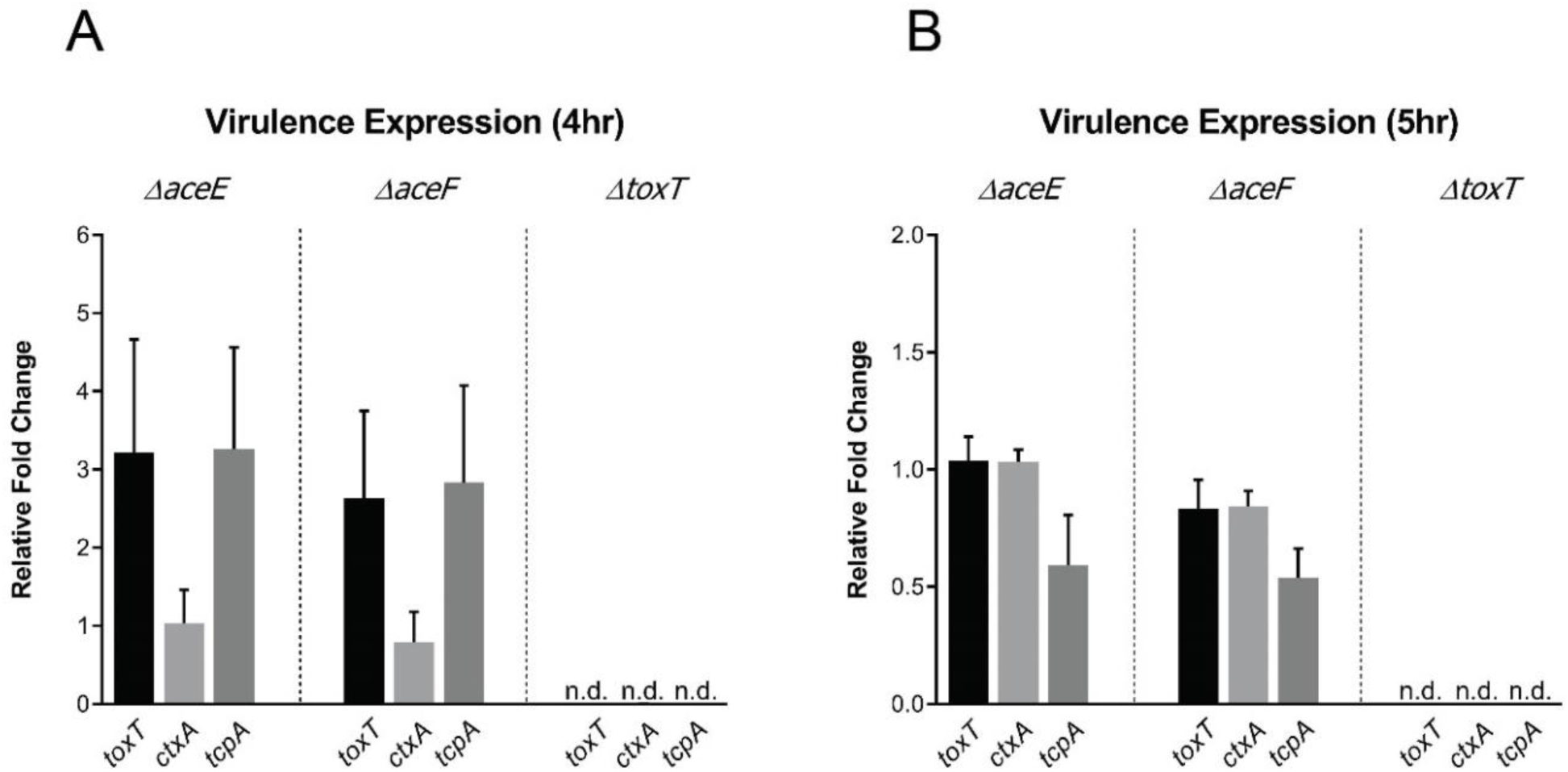
Relative fold change of *toxT, ctxA*, and *tcpA* transcript levels compared to wild type expression. (A-B) RNA was isolated from WT, *ΔaceE, ΔaceF*, and *ΔtoxT* cultures grown in standard AKI toxin-inducing conditions at 4h and 5h. Expression data was calculated by ΔΔCt using *recA* as an internal control. Data represent the average and SEM for three independent biological replicates.

### A functional pyruvate dehydrogenase complex is necessary for colonization of the infant mouse

Based on our findings that *V. cholerae* PDH mutants are defective for aerobic carbohydrate metabolism, we sought to examine whether or not this pathway was required to support growth *in vivo*. The infant mouse model is used extensively to investigate intestinal colonization by *V. cholerae* (49). Infant mice produce a mucus layer in the intestine that can serve as a substrate for *V. cholerae* growth (13), although they also have reduced resident microbiota and a less developed immune system compared to adult mice (50). The reduction in resident flora increases the necessity for *V. cholerae* to liberate mucin glycans for substrate utilization as the commensal population does not provide this resource as in other systems (51).

Here, we oro-gastrically infected CD-1 mouse neonates with ∼10^6^ CFU bacterial cells to compare colonization of wild type and PDH mutant *V. cholerae*. In mono-associated infections, PDH mutants were attenuated for colonization by approximately 100-fold compared to wild type (Figure 5A). We also assessed direct *in vivo* competition with wild type by co-infecting each mutant with a PDH+ *ΔlacZ* (VC2338) strain of *V. cholerae*. PDH mutant strains exhibited a similar attenuation in competition with wild type as was observed in mono-associated infections, with mutant recovery approximately 100-fold lower than wild type after co-infection (Figure 5B). These data indicate a requirement for a functionally active PDH complex to support colonization of the infant mouse, suggesting that oxidative metabolism of carbohydrate substrates is critical for colonization and population expansion.

**Figure 5.**
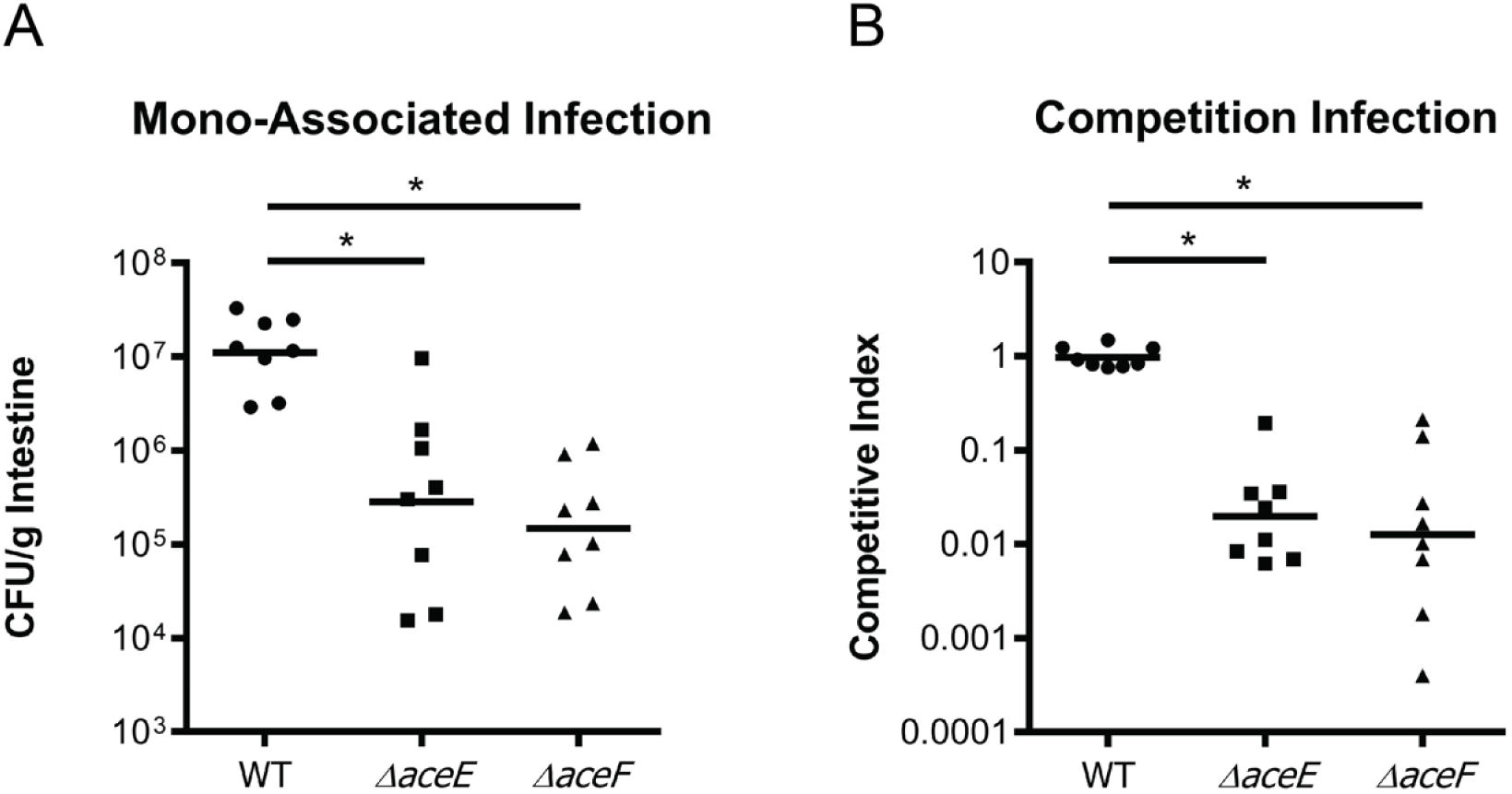
Infant mouse colonization assays of WT, *ΔaceE*, and *ΔaceF* after 20h. (A) Mono-associated infections of 3-5 day old infant mice reported as CFU/g intestine. (B) Competition infections of 3-5 day old infant mice reported as a competitive index score calculated as a ratio of output versus input [(Target_Output_/*ΔlacZ*_Output_) / (Target_Input_/*ΔlacZ*_Input_)]. WT, *ΔaceE*, and *ΔaceF* strains were co-inoculated with an *aceE+/*aceF+ *ΔlacZ* strain (PDH+) to determine the relative fitness of each test strain. Data for each experiment was obtained from 8 independent mouse colonization infections. The bar represents geometric mean. Statistical analysis was performed using GraphPad PRISM where significance was tested on log transformed data by ANOVA with post-hoc Tukey’s Test; * indicates p<0.05.

As oxygen levels (52, 53) and mucin composition (54) fluctuate along the length of the small intestine, we investigated the relative importance of PDH function across the longitudinal axis in the infant mouse intestine. 1cm long pieces of intestine were harvested from proximal, medial, and distal regions of the small intestine and assayed for recoverable CFU. Throughout the small intestine, *V. cholerae* PDH mutants were detected at levels well below that of wild type and in some cases were not detected as counts were below our limit of detection (Figure 6A-C). Here we again conclude that the PDH complex promotes *V. cholerae* colonization, and that oxidative metabolism of carbohydrates is a key feature of *V. cholerae* growth and proliferation along the entire length of the small intestine. Additionally, bacterial loads of the PDH mutants across the individual intestinal segments do not appear to reflect the CFU/g counts obtained from analyzing the entire gastrointestinal tract in the previous mono-associated infection (Figure 5A). This suggests that the majority of PDH mutants detected in the previous mono-infection experiment resided within either the cecum or large intestine, sites anticipated to support more anaerobic metabolism (53). This finding further supports the necessity for an active PDH particularly at the primary site of infection in the small intestine.

**Figure 6.**
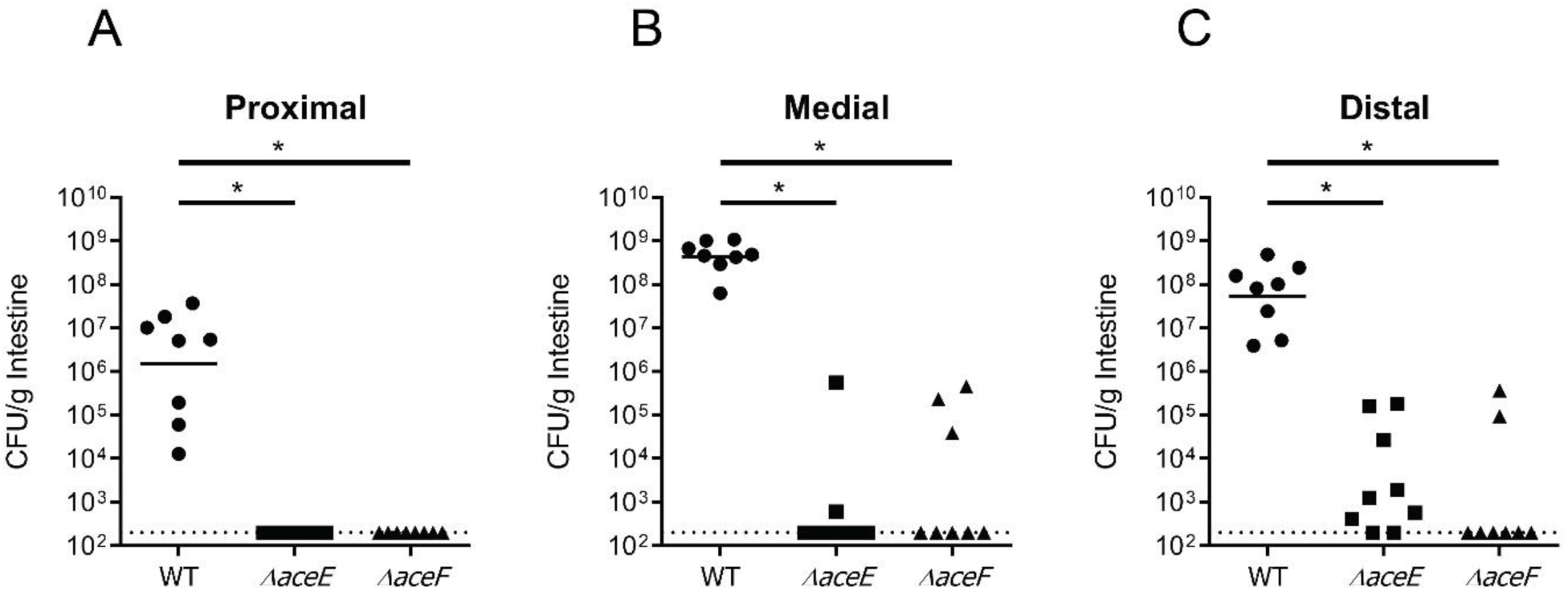
Infant mouse colonization of WT, *ΔaceE*, and *ΔaceF* mono-associated infections in (A) proximal, (B) medial, and (C) distal portions of the small intestine after 20h. Data for each segment was obtained from 8 independent mouse colonization infections. The bar represents the geometric mean and can only be plotted for the WT strain. Statistical analysis was performed using GraphPad PRISM where significance was tested on non-transformed data by Kruskal-Wallis analysis with post-hoc Dunn’s test; * indicates p<0.05.

### Pyruvate formate-lyase provides minor growth support during infection

As aerobic metabolism was determined to be beneficial to population expansion of *V. cholerae*, we wanted to explore the contributing effects of anaerobic metabolism to colonization. Oxygen gradation within the small intestine maintains highest oxygen availability in the intestinal crypts and nears hypoxia at the villus tip (28, 55). To determine if anaerobic proliferation also contributes to population expansion of *V. cholerae* during infection, potentially in the more anoxic lumen of the small intestine, CD-1 mice were infected with ∼10^6^ CFU of PFL mutant *ΔpflA*. To accurately assess the importance of PFL during infection, recovery of *V. cholerae* was performed for the small intestine separate from the large intestine. As the large bowel is inherently more anoxic, where PFL would be expected to function more readily, we focused more on assessing the role of PFL in the small intestine, as a more clinically relevant site for human *V. cholerae* infection.

In mono-associated infections, the PFL mutant was attenuated for colonization by approximately two-fold compared to wild type (Figure 7A). These data suggest that PFL, and therefore anaerobic metabolism, provides a less critical level of energy production than PDH to support growth during infection. Our results are consistent with previous reports that investigated anaerobic nitrate respiration demonstrating a similar 2-fold reduction in colonization of the infant mouse (56). However, in a competition experiment, the *ΔpflA* mutant colonized to levels equivalent to wild type (Figure 7B), essentially demonstrating no colonization defect at all. The same colonization pattern of the PFL mutant strain was observed in the large intestine (Supplemental Figure S8).

**Figure 7.**
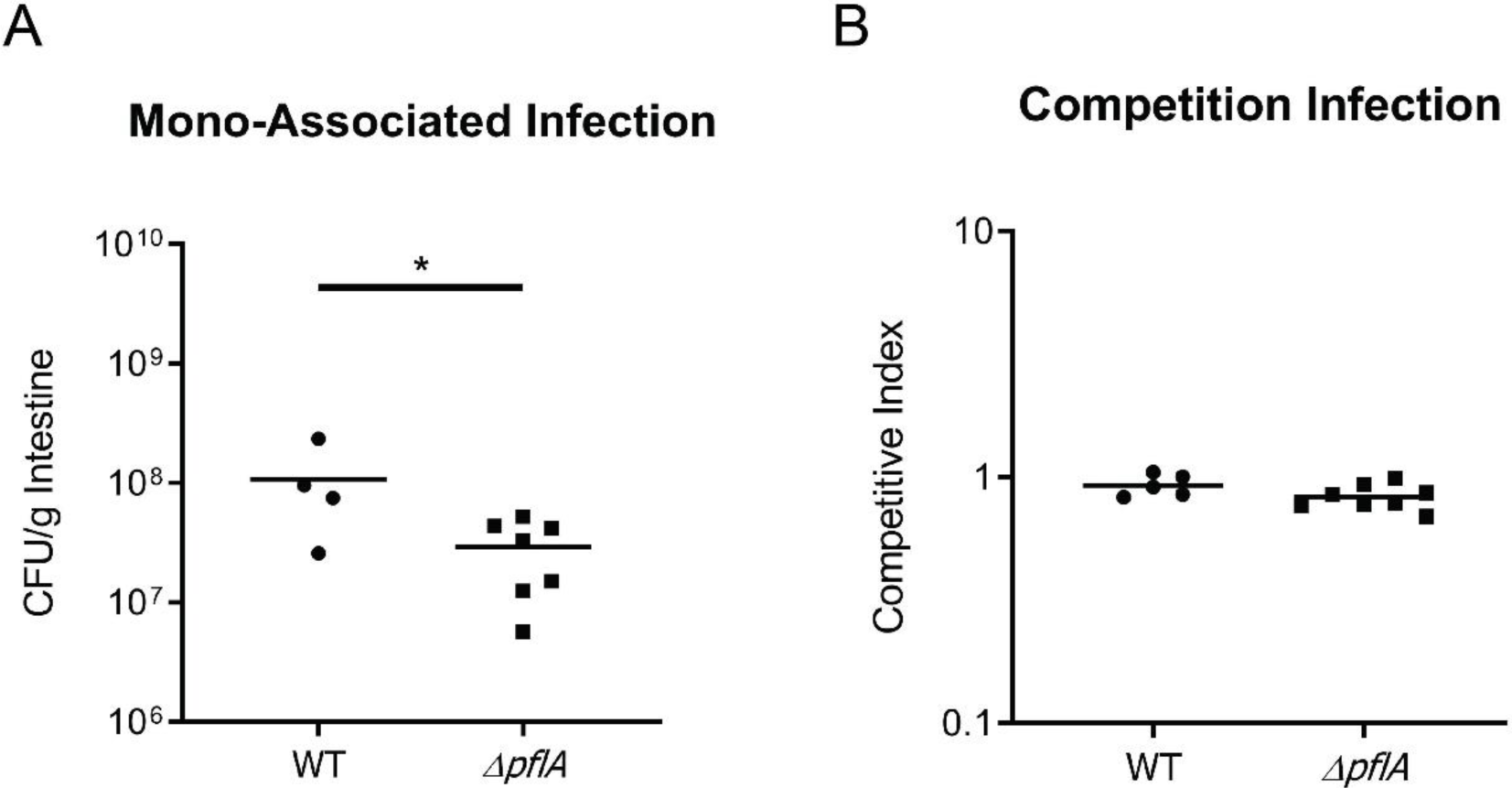
Infant mouse colonization assays of WT and *ΔpflA* in the small intestine after 20h. (A) Mono-associated infections of 3-5 day old infant mice reported as CFU/g intestine. (B) Competition infections of 3-5 day old infant mice reported as a competitive index score calculated as a ratio of output versus input [(Target_Output_/*ΔlacZ*_Output_) / (Target_Input_/*ΔlacZ*_Input_)]. WT and *ΔpflA* strains were co-inoculated with a *pflA*+ *ΔlacZ* strain to determine the relative fitness of each test strain. Data for each experiment was obtained from 4-5 independent mouse colonization infections for WT and 7-8 mouse infections for *ΔpflA*. The bar represents geometric mean. Statistical analysis was performed using Graphpad PRISM where significance was tested on log transformed data by Student’s t-test; * indicates p<0.05.

One hypothesis to explain this lack of fitness defect when co-infected with wild type is during co-infection, the *pflA+ ΔlacZ* strain may produce acetate that can be metabolized by *ΔpflA* to mitigate the two-fold defect seen in mono-associated infections (57). Acetate would provide acetyl-CoA by way of acetyl-CoA synthase-1 (ACS-1), circumventing the PDH/PFL carbohydrate utilization pathways (58). To test this hypothesis, we first demonstrated that all strains are capable of growth on acetate (Supplemental Figure S9). Then, to test whether metabolic rescue of the *ΔpflA* strain by *pflA+ ΔlacZ* occurs, monocultures and competition cultures were grown anaerobically in M9 minimal media with 0.5% glucose. At both the 12h and 20h timepoints, PFL mutant was found to be between 2-10 fold reduced compared to *pflA*+ *ΔlacZ* for both monoculture and competition comparisons. These findings indicate that the similar output ratios detected in competition *in vivo* assays are unlikely related to acetate supplementation by the *pflA*+*ΔlacZ* strain (Supplemental Figure S10).

## Discussion

Oxygen-dependent metabolism is key to pathogenicity in many gastrointestinal microbes. Pathogens that actively manipulate the host environment to oxygenate the gut rapidly proliferate during infection. Although a direct link has yet to be determined, the cholera toxin of *V. cholerae* may increase oxygen availability in the gut through its influence on optimal TCA cycle activity during infection (59). In other gastrointestinal pathogens, oxidative metabolism is supported by inducing inflammation at the site of infection. Inflammation in response to *Citrobacter rodentium* or *Salmonella* Typhimurium infections promotes colonization, proliferation of the microbe, and disease as a result of increased aerobic metabolism (60, 61). While *V. cholerae* infection does not lead to significant changes in gross pathology of intestinal architecture and cholera is not typically characterized as a pro-inflammatory infection, inflammatory markers are increased in animal models and human infection (62, 63). Whether oxygen levels in the gut are elevated due to this innate immune response has yet to be determined. There is evidence to support inflammation promoting *V. cholerae* colonization in some circumstances. In *V. cholerae* strain V52, the Type VI secretion system (T6SS) increases intestinal inflammation in the infant mouse and promotes increased colonization levels (64). Also, the newly emerged El Tor Haitian variant strain has higher virulence in animal models reaching a higher bacterial cell burden than previously characterized strains and causing elevated inflammation and epithelial cell damage (65). Evolved *V. cholerae* strains equipped to withstand an inflamed environment could benefit from increased oxygen availability to generate more energy to support growth and proliferation.

Oxygen contribution to *V. cholerae* pathogenicity has been examined principally in regard to the ToxR/TcpP/ToxT virulence cascade. For example, the regulators AphB and OhrR respond to reduced environmental oxygen by activating *tcpP* expression (66). Further, decreased oxygen levels in stationary culture conditions stimulates ToxR-TcpP interaction (67) and activation of *toxT* transcription (48). Translating these *in vitro* results to the context of oxygen distribution *in vivo* would imply that the anoxic lumen primes *V. cholerae* for virulence gene expression prior to accessing the more oxygenated host epithelium. The radial oxygen gradient within the intestine would therefore influence optimal timing of *V. cholerae* virulence expression. In this work, we sought to explore how oxygen-dependent and independent metabolic pathways influence population expansion during infection, but not necessarily as they relate to virulence gene expression.

Oxygen availability in the crypt spaces of the intestine, along with the presence of carbohydrate-rich mucin molecules could vastly improve growth and proliferation of *V. cholerae* in this site. The mucus lining of the gastrointestinal tract protects the host epithelium from both resident and transient microorganisms to maintain gut homeostasis (68). *V. cholerae* has the capacity to bypass this host defense through motility (69) as well as to exploit it for growth substrates. However, this does not suggest that the mucus layer is inconsequential to curtailing the effects of *V. cholerae* pathogenicity. In mice with a chemically degraded mucus layer, *V. cholerae* bacterial counts exceeded that of untreated mice (13) indicating that mucus contributes to abatement of disease. Similarly, *Muc2*^*-/-*^ mice that lack the primary secretory mucin of the intestinal tract, MUC2, exhibit inflammation in the large intestine due to commensal population interactions with the epithelium as well as exacerbated infection in *Citrobacter rodentium* and *Salmonella* Typhimurium challenge models (70–72). Thus, the mucus serves a protective function against *V. cholerae* yet is exploited to benefit the microbe.

The sequential progression of *V. cholerae* pathogenicity is tightly linked to bacterial-mucin interactions. Adherence to host mucin by GbpA, an *N*-acetylglucosamine binding protein, is a key step in colonizing the small intestine (73). Upon reaching the host epithelium *V. cholerae* establishes an adherent microcolony (74). Within this microenvironment, mucin breakdown products and the presence of oxygen help drive the population expansion of *V. cholerae* (this work). Stimulation of the host epithelium by cholera toxin induces production and secretion of goblet cell mucin (75) in addition to other host-derived nutrients such as iron and long-chain fatty acids (59), and potentially oxygen. As the population of cells rapidly expands, mucin breakdown products stimulate motility of *V. cholerae* (76), a trait required for optimal colonization of the proximal and medial portions of the small intestine (13, 77, 78). This interaction likely contributes to population dynamics observed for *V. cholerae* whereby it migrates counter to intestinal flow in the later stages of infection to populate the proximal and medial portions of the intestine (79). This motility response acts in coordination with the secreted mucolytic hemagglutinin/protease (HapA) of *V. cholerae*, used during cellular detachment (80, 81). HapA is stimulated both by high cell density through the activity of the regulator HapR, and directly when in the presence of mucin (82). As *V. cholerae* exits the host, a fraction of the population is embedded in mucin (83) which may influence hyper-infectivity of human-passaged *V. cholerae* (84). From initial inoculation into the human gut to the eventual passaging of the bacteria, interactions between *V. cholerae* and mucus substantially influence *V. cholerae* pathogenicity.

Our work postulates that mucin metabolism enhances proliferation of *V. cholerae* during the course of infection. *V. cholerae* is a facultative anaerobe, and we sought to uncover whether aerobic or anaerobic metabolism enables it to grow to high levels during infection. We assessed the *in vivo* fitness of strains lacking either the *aceE* or *aceF*-encoded components of the pyruvate dehydrogenase (PDH) complex, or *pflA-*encoded pyruvate formate-lyase (PFL). These enzymes catalyze production of acetyl-CoA from pyruvate either aerobically (PDH) or anaerobically (PFL). In *Escherichia coli*, PFL is induced only during anaerobiosis whereas the PDH complex can function in both anaerobic and anaerobic environments (85, 86). However, unlike what is observed in *E. coli*, in our work, *V. cholerae* lacking PFL was not rescued for anaerobic growth to any noticeable extent by having a functional PDH. This enabled us to differentiate the contribution of aerobic and anaerobic metabolism by investigating PDH and PFL mutants. The significant loss of fitness by the *ΔaceE*/*ΔaceF* strains compared to the wild type strain suggests that *V. cholerae* population expansion in the small intestine is driven largely by aerobic, oxidative metabolism. This is consistent with *V. cholerae* preferentially localizing to the epithelial crypts (13), with greater oxygenation that enables oxidative metabolic pathways to generate energy (28). The radial distribution of oxygen in the intestine therefore biogeographically relegates replicative *V. cholerae* cells primarily to the epithelium as opposed to the more anoxic lumen. Our results do not completely rule out some anaerobic growth and expansion of *V. cholerae* during infection as the *ΔpflA* mutant colonized to levels about half those of wild type, similar to what is observed with other anaerobic metabolism deficient strains (25, 44, 56). Additionally, there is recent evidence to suggest that multiple anaerobic metabolic pathways function in tandem to support growth in anoxic conditions. A double mutant in ethanol fermentation and nitrate respiration showed a significant reduction in colonization compared with single mutants that were near wild type levels of colonization (87).

While expansion of *V. cholerae in vivo* evidently proceeds primarily through aerobic production of acetyl-CoA using PDH as opposed to anaerobic acetyl-CoA production using PFL, how it uses its reducing equivalents to generate energy from the electron transport chain is less certain from our work. Oxygen as a terminal electron acceptor is certainly possible given the availability of oxygen within the crypt epithelium and the presence of four terminal oxidase complexes in the *V. cholerae* genome (88), which are the subject of future investigation. *V. cholerae* also maintains nitrate, fumarate, TMAO, and DMSO reductases that can function as terminal electron acceptors in anaerobic respiration (88). Fumarate and TMAO support *V. cholerae* growth in anaerobic conditions (44) as does nitrate when in an alkaline environment (56), albeit not to the extent observed for oxidative growth. This is primarily due to the relatively low redox potentials of fumarate and TMAO relative to O_2_ and *V. cholerae* requiring alkaline pH environments for nitrate respiration as it lacks a nitrite reductase needed to eliminate this toxic compound (56). DMSO on the other hand was not shown to support *V. cholerae* growth at all (44). However, growth *in vivo* with addition of the alternative electron acceptor TMAO induces high levels of cholera toxin (44). Infant mice infected with an inoculum of El Tor strain N16961 mixed with TMAO exhibited more severe signs of infection, suggesting a TMAO-dependent toxin production effect. Although anaerobic terminal reductases may not be the principal mode of *V. cholerae* growth and expansion *in vivo*, they are still likely to contribute to *V. cholerae* pathogenesis. Resolution of the role of different terminal reductases regarding growth and pathogenicity *in vivo* also await future examination by investigating *V. cholerae* terminal reductase mutants.

