## Supplementary material for "Aerobic metabolism in *Vibrio cholerae* is required for population expansion during infection": Supplmental Material

Supplementary Table 1. Transposon mutagenesis screen results.

| NR Well Location | Locus Tag | Gene Name | Gene Description | Gene Role |
|---|---|---|---|---|
| 9-E1 | VC2482 | ilvH | acetolactate synthase III, small subunit | Amino acid biosynthesis |
| 10-D1 | VC0029 | ilvE | branched-chain amino acid amiotransferase | Amino acid biosynthesis |
| 9-E12 | VCA0684 | thiE | thiamin-phosphate pyrophosphorylase | Biosynthesis of cofactors, prosthetic groups, and carriers |
| 17-F8 | VC0061 | thiC | thiamin biosynthesis protein ThiC | Biosynthesis of cofactors, prosthetic groups, and carriers |
| 33-G10 | VC1296 | thiD | phosphomethylpyrimidine kinase | Biosynthesis of cofactors, prosthetic groups, and carriers |
| 12-E7 | VC2481 | serA | D-3-phosphoglycerate dehydrogenase | Amino acid biosynthesis |
| 28-B11 | VC2345 | serB | phosphoserine phosphatase | Amino acid biosynthesis |
| 28-H7 | VC2363 | thrB | homoserine kinase | Amino acid biosynthesis |
| 33-G6 | VC2362 | thrC | threonine synthase | Amino acid biosynthesis |
| 10-F1 | VC1169 | trpA | tryptophan synthase, alpha subunit | Amino acid biosynthesis |
| 33-H10 | VC1170 | trpB | tryptophan synthase, beta subunit | Amino acid biosynthesis |
| 34-C5 | VC1174 | trpE | anthranilate synthase component I | Amino acid biosynthesis |
| 5-H6 | VC0705 | pheA | chorismate mutase/prephenate dehydratase | Amino acid biosynthesis |
| 25-C3 | VC0056 | aroE | shikimate 5-dehydrogenase | Amino acid biosynthesis |
| 9-B1 | VC0149 | epsC | general secretion pathway protein C | Protein fate |
| 28-E3 | VC2725 | epsL | general secretion pathway protein L | Protein fate |
| 33-E3 | VC1709 |  | zinc protease, insulinase family | Protein fate |
| 1-F3 | VC1181 | cydD | transport ATP-binding protein CydD | Transport and binding proteins |
| 12-G7 | VC2270 | ribE | riboflavin synthase, alpha subunit | Biosynthesis of cofactors, prosthetic groups, and carriers |
| 23-C6 | VC2268 | ribE | 6,7-dimethyl-8-ribityllumazine synthase | Biosynthesis of cofactors, prosthetic groups, and carriers |
| 28-H2 | VC0094 | ubiA | 4-hydroxybenzoate octaprenyltransferase | Biosynthesis of cofactors, prosthetic groups, and carriers |
| 23-C8 | VC2628 | aroB | 3-dehydroquinate synthase | Amino acid biosynthesis |
| 5-A12 | VC2414 | aceE | pyruvate dehydrogenase, E1 component | Energy metabolism |
| 32-A1 | VC2413 | aceF | pyruvate dehydrogenase, E2 component, dihydrolipoamide acetyltransferase | Energy metabolism |
| 27-A7 | VC2544 | fbp | fructose-1,6-bisphosphatase | Energy metabolism |
| 4-H4 | VC0604 | acnB | aconitate hydratase 2 | Energy metabolism |
| 26-A10 | VC1487 |  | conserved hypothetical protein | Hypothetical proteins |
| 3-C8 | VC0849 |  | conserved hypothetical protein | Hypothetical proteins |

Supplementary Table 2. Bacteria strain list.

| **Strain** | **Description** | **Reference** |
|---|---|---|
| ***Vibrio cholerae*** | | |
| *V. cholerae* C6706 El Tor biotype (Wild type) | Wild type strain | Waters Lab Collection |
| *V. cholerae ΔaceE* (VC2414) | Isogenic deletion strain | This study |
| *V. cholerae ΔaceF* (VC2413) | Isogenic deletion strain | This study |
| *V. cholerae ΔpflA* (VC1869) | Isogenic deletion strain | This study |
| *V. cholerae ΔlacZ* (VC2338) | Isogenic deletion strain | This study |
| *V. cholerae ΔtoxT* (VC0838) | Isogenic deletion strain | Waters Lab Collection |
| *V. cholerae* pMMB66EH (empty vector) | Complementation verification strain | This study |
| *V. cholerae ΔaceE* pMMB66EH (empty vector) | Complementation verification strain | This study |
| *V. cholerae ΔaceE* pMMB66EH-*aceE* | Complementation verification strain | This study |
| *V. cholerae ΔaceF* pMMB66EH (empty vector) | Complementation verification strain | This study |
| *V. cholerae ΔaceF* pMMB66EH-*aceF* | Complementation verification strain | This study |
| *V. cholerae ΔpflA* pMMB66EH (empty vector) | Complementation verification strain | This study |
| *V. cholerae ΔpflA* pMMB66EH-*pflA* | Complementation verification strain | This study |
| ***Escherichia coli*** | | |
| *E. coli* MCH100λpir pKAS32 (empty vector) | Plasmid vector strain | Lab Collection |
| *E. coli* S17 pMMB66EH (empty vector) | Plasmid vector strain | Lab Collection |
| *E. coli* S17 λpir 3-7 | Cloning vector recipient | Lab Collection |

### Supplementary Table 3. Primer list.

| | Primer Name | Primer Sequence (5' -> 3') | Description | Reference |
|---|---|---|---|---|
| **Mutant Construction Primers** | | | | |
| *aceE* VC2414 | *aceE* Upstream Homology F | GTGGAATTCCCGGGAGAGCTCAATATTTTTGCTGTTTAATCAACTCTTG | pKAS32 construction primer | This study |
| | *aceE* Upstream Homology R | CATTACTTTTCCTACCTTCAAGGCGATCTATCCTTCTGTTGG | pKAS32 construction primer | This study |
| | *aceE* Downstream Homology F | CCAACAGAAGGATAGATCGCCTTGAAGGTAGGAAAAGTAATG | pKAS32 construction primer | This study |
| | *aceE* Downstream Homology R | CCGCGGACATGTACAGAGCTGCACGACTGGAGAAGCATG | pKAS32 construction primer | This study |
| | *aceE* pKAS32 Seq Primer F1 | GAACTGGAGAGACTGATAGTGGAAG | pKAS32 sequencing primer | This study |
| | *aceE* pKAS32 Seq Primer F2 | CAGAATTTTAAACTCTTACATCGC | pKAS32 sequencing primer | This study |
| | *aceE* pKAS32 Seq Primer R1 | CAAAAGGTGCAGGTACTTCCAT | pKAS32 sequencing primer | This study |
| | *aceE* pKAS32 Seq Primer R2 | CTGCACCTTCCGCTTCAA | pKAS32 sequencing primer | This study |
| | *aceE* Deletion Detection F | CACCTCTAGCCCATCAAGTCC | Isogenic deletion verification primer | This study |
| | *aceE* Deletion Detection R | GTAGTTCTGTACGTCTTCTTTCAGG | Isogenic deletion verification primer | This study |
| *aceF* VC2413 | *aceF* Upstream Homology F | GTGGAATTCCCGGGAGAGCTTTCTTACTACAAAGAAGCGACTTC | pKAS32 construction primer | This study |
| | *aceF* Upstream Homology R | TTTCGCCACCCGAGAATGCGCTTAAGCGTACAGCGGGTTGGT | pKAS32 construction primer | This study |
| | *aceF* Downstream Homology F | ACCAACCCGCTGTACGCTTAAGCGCATTCTCGGGTGGCGAAA | pKAS32 construction primer | This study |
| | *aceF* Downstream Homology R | CCGCGGACATGTACAGAGCTCTTTCGCTTCAACGGCGGTC | pKAS32 construction primer | This study |
| | *aceF* pKAS32 Seq Primer F1 | GTACAACGCTGAACGGTGAA | pKAS32 sequencing primer | This study |
| | *aceF* pKAS32 Seq Primer F2 | ATCGCAGCGACTGACTACAT | pKAS32 sequencing primer | This study |
| | *aceF* pKAS32 Seq Primer R1 | CAGCGCATCGGTTGAATC | pKAS32 sequencing primer | This study |
| | *aceF* pKAS32 Seq Primer R2 | GCTCATTTCGACCTCTTGTAGTC | pKAS32 sequencing primer | This study |
| | *aceF* Deletion Detection F | TCCGTCAAATCGGTATCTACA | Isogenic deletion verification primer | This study |
| | *aceF* Deletion Detection R | CGCATCGTAACGCTCAGCT | Isogenic deletion verification primer | This study |
| *pflA* VC1869 | *pflA* Upstream Homology F | GTGGAATTCCCGGGAGAGCTTCACTGACTCGCAAAAAAG | pKAS32 construction primer | This study |
| | *pflA* Upstream Homology R | AGAGGATGAAGAGCGCTTCTCAGTTATG | pKAS32 construction primer | This study |
| | *pflA* Downstream Homology F | AGAAGCGCTCTTCATCCTCTCGACGTTATC | pKAS32 construction primer | This study |
| | *pflA* Downstream Homology R | TGCGCATGCTAGCTATAGTTAACTCGCGGTTCAGTTCAC | pKAS32 construction primer | This study |
| | *pflA* pKAS32 Seq Primer F | AATCTCAGACACCTTGTTTGAC | pKAS32 sequencing primer | This study |
| | *pflA* pKAS32 Seq Primer R | GCCAGATATAAAGGGAGTTAAGC | pKAS32 sequencing primer | This study |
| | *pflA* Deletion Detection F | GCTGTTGCTTCACTACTGAG | Isogenic deletion verification primer | This study |
| | *pflA* Deletion Detection R | GGATTCTTGCATCGCATGATAC | Isogenic deletion verification primer | This study |
| *lacZ* VC2338 | *lacZ* Upstream Homology F | GTGGAATTCCCGGGAGAGCTGCCACCAAACACTAAGCTTC | pKAS32 construction primer | This study |
| | *lacZ* Upstream Homology R | GCTCTCTGGCCCCTCAAGCCGAGGAGTAAAG | pKAS32 construction primer | This study |
| | *lacZ* Downstream Homology F | GGCTTGAGGGGCCAGAGAGCCTTAAGGC | pKAS32 construction primer | This study |
| | *lacZ* Downstream Homology R | TGCGCATGCTAGCTATAGTTTAGCACGTGAAGCCGGTG | pKAS32 construction primer | This study |
| | *lacZ* pKAS32 Seq Primer F | GATAACCAATCGCAAAACCAACTT | pKAS32 sequencing primer | This study |
| | *lacZ* pKAS32 Seq Primer R | TCTCATCCGTCAAGGACATAGAAAC | pKAS32 sequencing primer | This study |
| | *lacZ* Deletion Detection F | GAAATTGATCGGTCGATAGGCTG | Isogenic deletion verification primer | This study |
| | *lacZ* Deletion Detection R | CCGAGTCCATAACTCTTATCCTTCTTA | Isogenic deletion verification primer | This study |
| pKAS32 | pKAS32 Multiple Cloning Site Seq Primer F | GCCTCTAAGGTTTTAAGTTT | pKAS32 specific sequencing primer | Lab Collection |
| | pKAS32 Multiple Cloning Site Seq Primer R | CTTTCAAGGTAGCGGTTACC | pKAS32 specific sequencing primer | Lab Collection |
| *toxT* VC0838 | 1531 | CAACTTCTGTAGTTAATGCAATTCCC | toxT deletion verification primer | Waters Lab Collection |
| | 1532 | CCCTCCAGTAAATTTTCATAAATGTCG | toxT deletion verification primer | Waters Lab Collection |
| **Complementation Primer Sets** | | | | |
| *aceE* VC2414 | pMMB66EH *aceE* ORF F | CAGGAAACAGAATTCCCGGGATGTCTGACATGAAGCATGAC | pMMB66EH *aceE* ORF construct primer | This study |
| | pMMB66EH *aceE* ORF R | CTCATCCGCCAAAACAGCCATTAAGCGTACAGCGGGTGG | pMMB66EH *aceE* ORF construct primer | This study |
| | pMMB66EH *aceE* ORF Seq Primer 1 | CTGCGTGCATCGAAGAAAGA | pMMB66EH *aceE* ORF sequencing primer | This study |
| | pMMB66EH *aceE* ORF Seq Primer 2 | CTGTTATGGGTAACGGTAAG | pMMB66EH *aceE* ORF sequencing primer | This study |
| | pMMB66EH *aceE* ORF Seq Primer 3 | GTACCTGAAACTGGAAGAAG | pMMB66EH *aceE* ORF sequencing primer | This study |
| | pMMB66EH *aceE* ORF Seq Primer 4 | GCGTGGATGGCGGGTGAC | pMMB66EH *aceE* ORF sequencing primer | This study |
| *aceF* VC2413 | pMMB66EH *aceF* ORF F | CAGGAAACAGAATTCCCGGGTTGAAGGTAGGAAAAGTAATG | pMMB66EH *aceF* ORF construct primer | This study |
| | pMMB66EH *aceF* ORF R | CTCATCCGCCAAAACAGCCATTACAGTACCAGACGACG | pMMB66EH *aceF* ORF construct primer | This study |
| | pMMB66EH *aceF* ORF Seq Primer 1 | AGCAGCGGCAGCACCAGC | pMMB66EH *aceF* ORF sequencing primer | This study |
| | pMMB66EH *aceF* ORF Seq Primer 2 | AGATCAAAGTGGCTACAGGCGA | pMMB66EH *aceF* ORF sequencing primer | This study |
| | pMMB66EH *aceF* ORF Seq Primer 3 | GAGCAAAACGCGATGGAAGC | pMMB66EH *aceF* ORF sequencing primer | This study |
| *pflA* VC1869 | pMMB66EH *pflA* ORF F | CAGGAAACAGAATTCCCGGGATGTCTACCATTGGTCGAATTC | pMMB66EH *pflA* ORF construct primer | This study |
| | pMMB66EH *pflA* ORF R | CTCATCCGCCAAAACAGCCATCAATATTTTACGTTTGAGTGATAC | pMMB66EH *pflA* ORF construct primer | This study |
| pMMB66EH | pMMB66EH Multiple Cloning Site Seq Primer F | TGCATAATTCGTGTCGCTCA | pMMB66EH specific sequencing primer | This study |
| | pMMB66EH Multiple Cloning Site Seq Primer R | CTACGGCGTTTCACTTCTGA | pMMB66EH specific sequencing primer | This study |
| **RT-qPCR Primer Sets** | | | | |
| *toxT* VC0838 | toxT qPCR F | ACTGATGATCTTGATGCTATGGAG | qPCR Primer | This study |
| | toxT qPCR R | CATCCGATTCGTTCTTAATTCACC | qPCR Primer | This study |
| *ctxA* VC1475 | *ctxA* qPCR F | TGGAATCCCACCTAAAGCAG | qPCR Primer | This study |
| | *ctxA* qPCR R | TTGTTAGGCACGATGATGGA | qPCR Primer | This study |
| *recA* VC0543 | *recA* qPCR F | GGGTAACCTCAAGCAATCCA | qPCR Primer | This study |
| | *recA* qPCR R | CCACTCTTCGCCTTCTTTG | qPCR Primer | This study |
| *tcpA* VC0828 | *tcpA* qPCR F | ACGCAAATGCTGCTACACAG | qPCR Primer | This study |
| | *tcpA* qPCR R | CCCCTACGCTTGTAACCAAA | qPCR Primer | This study |

Supplementary Table 4. Purified porcine small intestinal mucin monosaccharide and sialic acid analysis determined by High-Performance Anion-Exchange Chromatography coupled with Pulsed Amperometric Detection (HPAEC-PAD). (GlycoAnalytics).

| **Monosaccharide** | **Amount (nmole/5µg of sample)** |
| --- | --- |
| Fucose | 0.17 |
| Galactosamine | 0.14 |
| Glucosamine | 0.14 |
| Galactose | 0.17 |
| Glucose | 0.01 |
| Mannose | 0.02 |

| **Sialic Acid** | **Amount (pmole/2µg of sample)** |
| --- | --- |
| Neu5Ac | 8.54 |
| Neu5Gc | 6.90 |

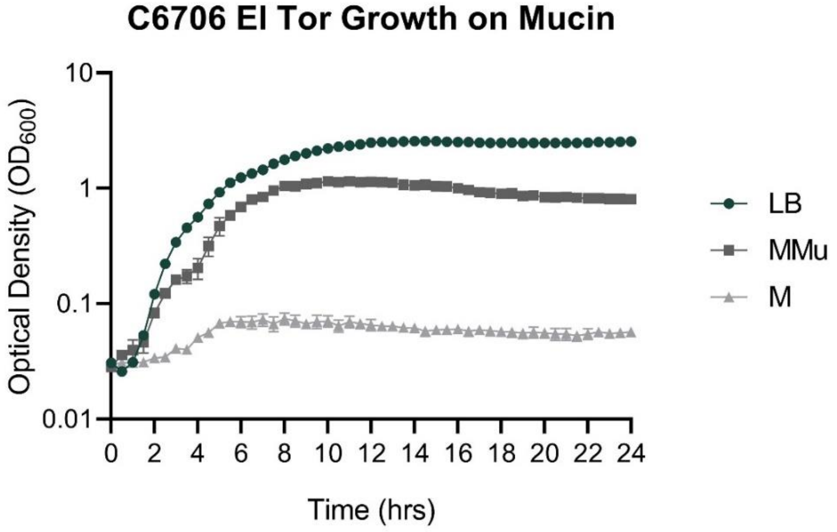


Supplementary Figure S1. *V. cholerae* C6706 El Tor wild type growth in LB, minimal 0.5% mucin (Sigma Type III porcine gastric mucin) (MMu), and minimal media with no added carbon source (M). Data represent the average and SEM for three independent biological replicates.

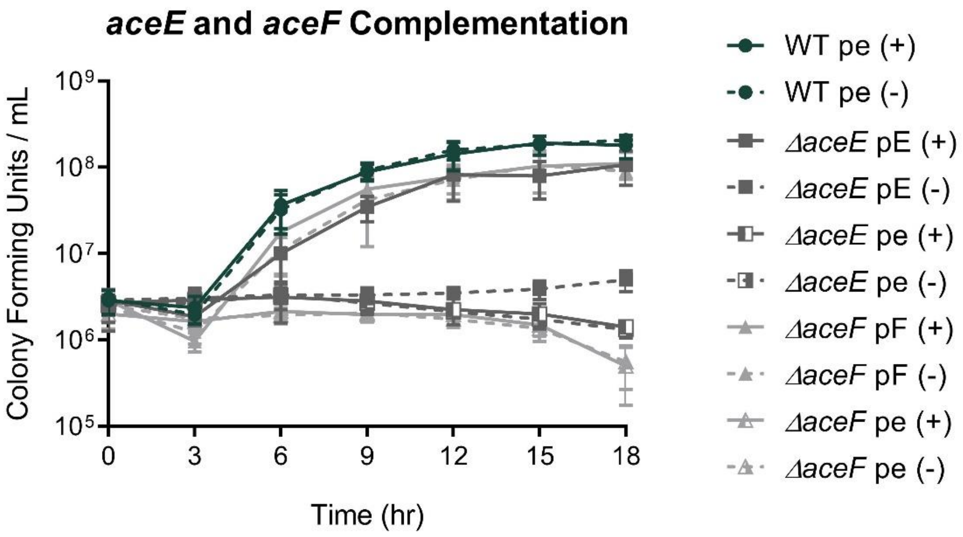


Supplementary Figure S2. Complementation growth curves of *ΔaceE* and *ΔaceF* in M9 0.5% glucose media. *ΔaceE* and *ΔaceF* strains were complemented with IPTG-inducible vector pMB66EH. WT and *ΔaceE* strains were induced with 1mM IPTG whereas *ΔaceF* was induced with 0.01mM IPTG. (+) indicates the addition of IPTG inducer whereas (-) indicates the lack thereof. 'pe' denotes an empty vector control whereas 'pE' and 'pF' indicate the complementation plasmid contains the *aceE* and *aceF* open reading frames, respectively. Data represent the average and SEM for three independent biological replicates.

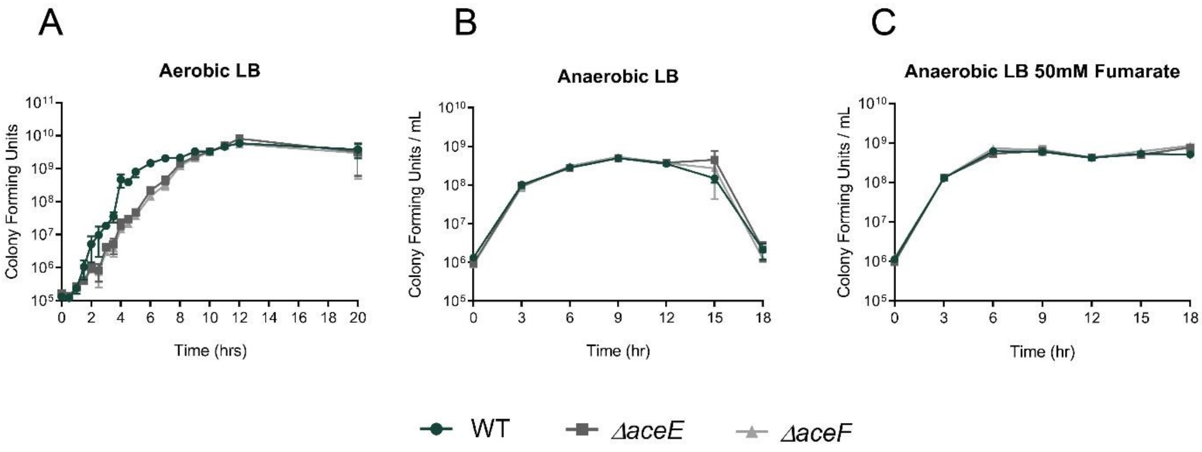


Supplementary Figure S3. Growth curves of WT (green circle) *ΔaceE* (dark grey square) and *ΔaceF* (grey triangle) in LB media grown (A) aerobically, (B) anaerobically, or (C) anaerobically supplemented with 50mM fumarate. Data represent the average and SEM for three independent biological replicates.

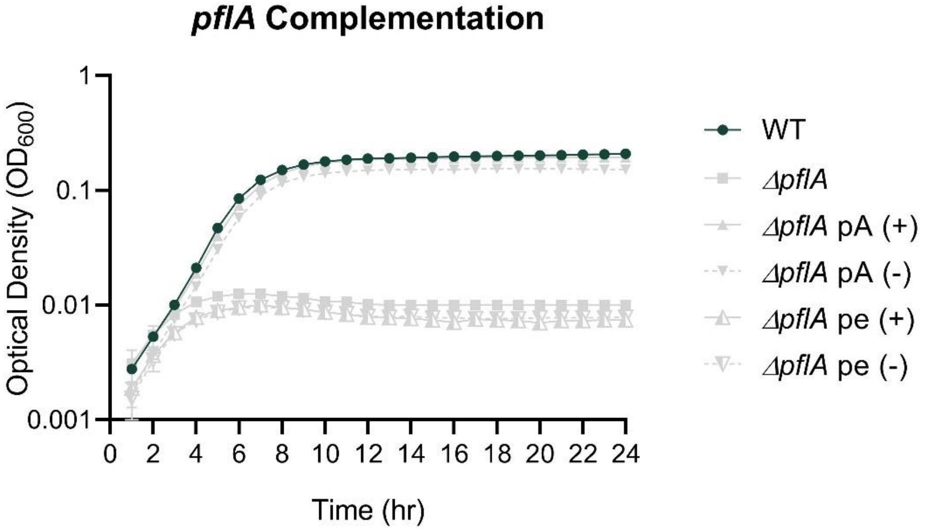


Supplementary Figure S4. Complementation growth curve of *ΔpflA* in M9 0.5% glucose 50mM fumarate media grown anaerobically. *ΔpflA* strain was complemented with IPTG-inducible vector pMB66EH. WT and *ΔpflA* strains were induced with 1mM IPTG. (+) indicates the addition of IPTG inducer whereas (-) indicates the lack thereof. ‘pe’ denotes an empty vector control whereas ‘pA’ indicates the complementation plasmid contains the *pflA* open reading frame. Data represent the average and SEM for three independent biological replicates.

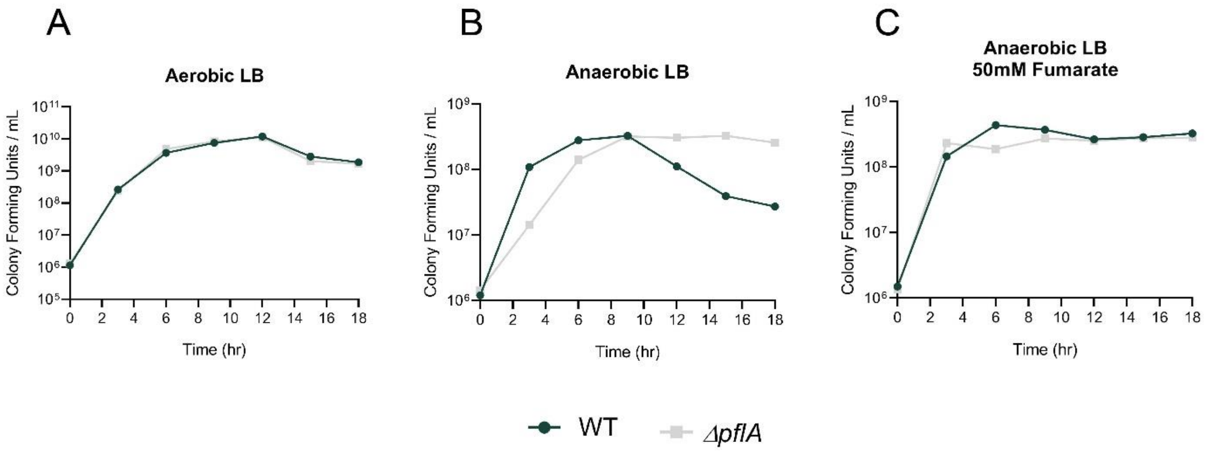


Supplementary Figure S5. Growth curves of WT (green circle) and *ΔpflA* (light grey square) in LB media grown (A) aerobically, (B) anaerobically, or (C) anaerobically supplemented with 50mM fumarate. Data represent the average and SEM for three independent biological replicates.

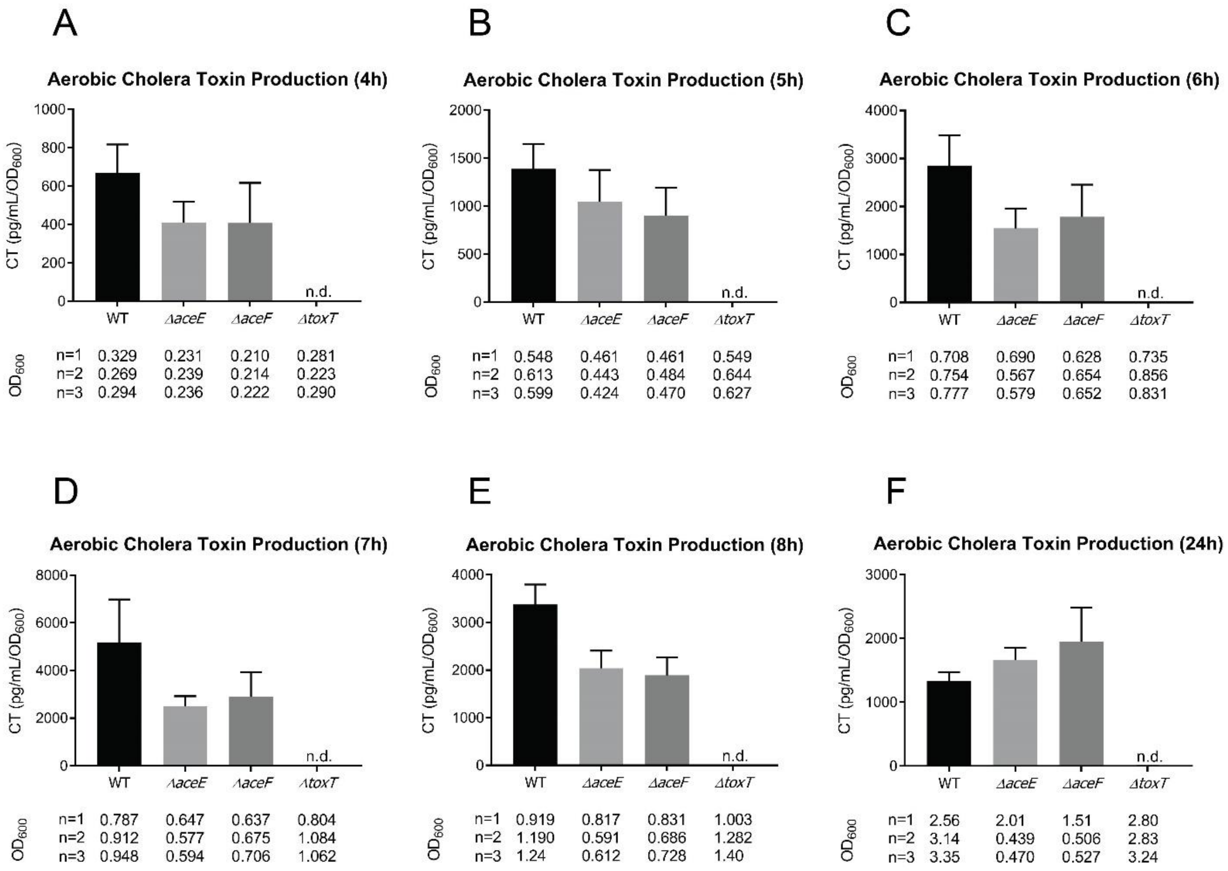


Supplementary Figure S6. Cholera toxin (CT) production for WT, *ΔaceE*, *ΔaceF*, and *ΔtoxT* strains. CT values relative to optical density (pg/mL/$OD_{600}$) are reported under standard AKI toxin-inducing conditions. (A-F) CT levels were measured at 4h, 5h, 6h, 7h, 8h, and 24h. The optical densities for the biological replicates are displayed below the corresponding strain on the x-axis. Data represent the average and SEM for three biological replicates.

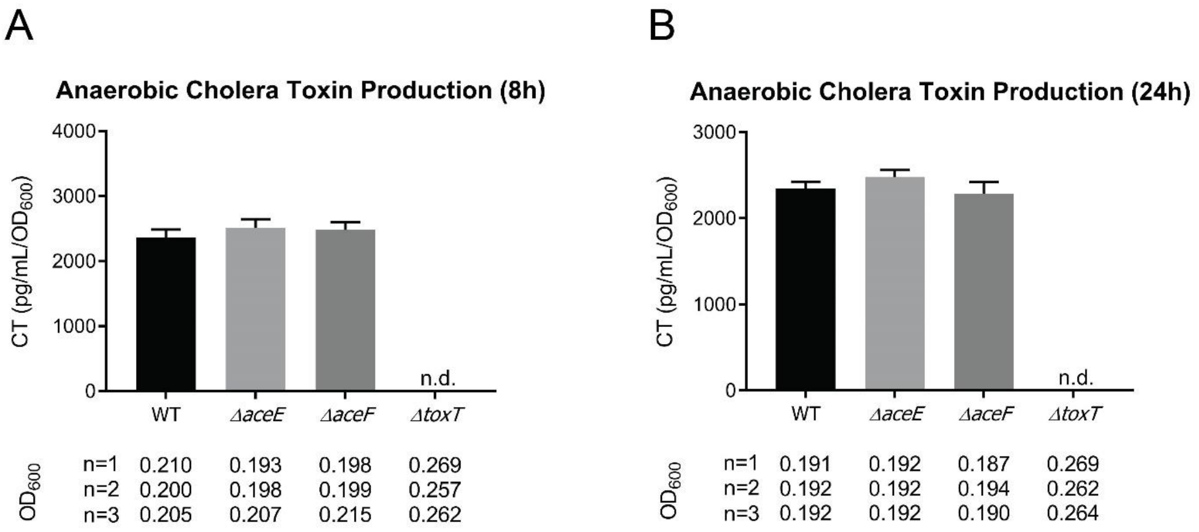


Supplementary Figure S7. Cholera toxin (CT) production for WT, *ΔaceE*, *ΔaceF*, and *ΔtoxT* strains. (A-B) CT values relative to optical density (pg/mL/$OD_{600}$) are reported under anaerobic AKI toxin-inducing conditions. CT levels were measured at 8h and 24h. The optical densities for the biological replicates are displayed below the corresponding strain on the x-axis. Data represent the average and SEM for three biological replicates. Panel A is a duplicate graph of Figure 3B and is included here for convenient comparison with the 24h timepoint.

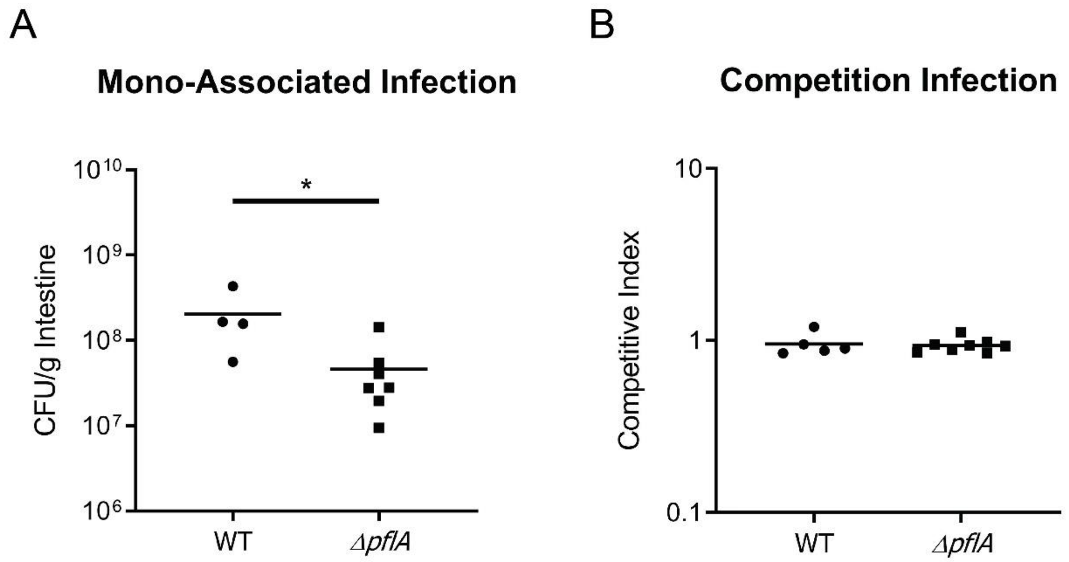


Supplementary Figure S8. Infant mouse colonization assays of WT and *ΔpflA* in the large intestine after 20h. (A) Mono-associated infections of 3-5 day old infant mice reported as CFU/g intestine. (B) Competition infections of 3-5 day old infant mice reported as a competitive index score calculated as a ratio of output versus input [(Target$_{Output}$/*ΔlacZ*$_{Output}$) / (Target$_{Input}$/*ΔlacZ*$_{Input}$)]. WT and *ΔpflA* strains were co-inoculated with a *pflA*+ *ΔlacZ* strain to determine the relative fitness of each test strain. Data for each experiment was obtained from 4-5 independent mouse colonization infections for WT and 7-8 mouse infections for *ΔpflA*. The bar represents geometric mean. Statistical analysis was performed using GraphPad PRISM where significance was tested on log transformed data by Student's t-test; * indicates $p<0.05$.

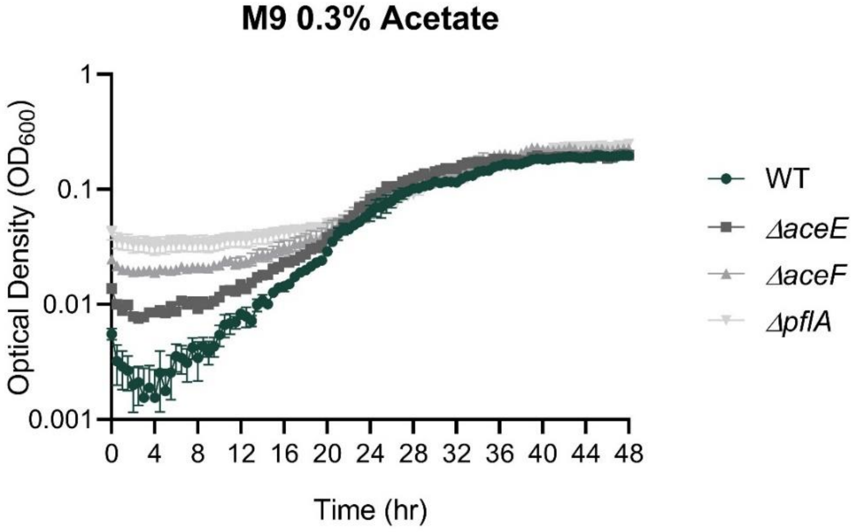


Supplementary Figure S9. Growth curves of WT (green circle), *ΔaceE* (dark grey square), *ΔaceF* (grey triangle) and *ΔpflA* (light grey inverted triangle) in M9 0.3% acetate. Data represent the average and SEM for three independent biological replicates.

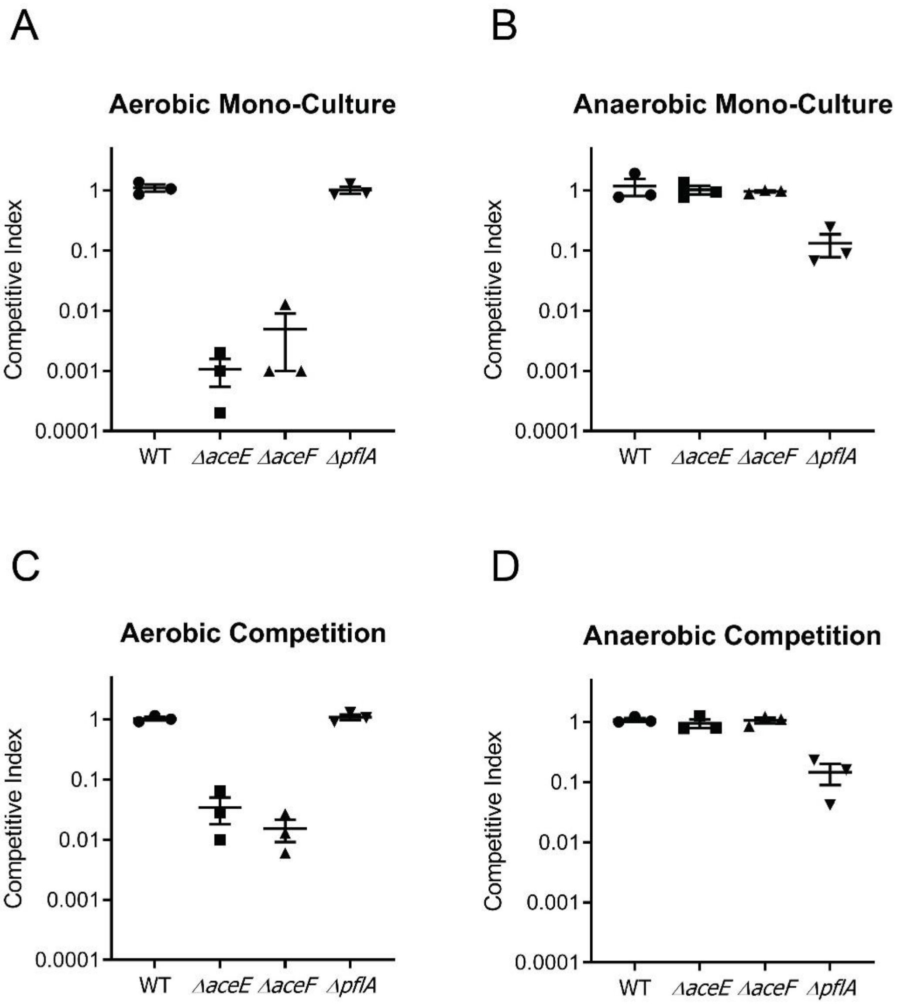


Supplementary Figure S10. *In vitro* mono-culture and competition assays of WT, *ΔaceE*, *ΔaceF*, and *ΔpflA* in M9 0.5% glucose after 20h. (A-B) Mono-culture competitive index scores were calculated as a ratio of endpoint culture to initial culture of test strains versus a PDH+/PFL+ *ΔlacZ* strain grown in separate culture tubes; $[(\text{Target}_{\text{Endpoint}}/\Delta lacZ_{\text{Endpoint}}) / (\text{Target}_{\text{Initial}}/\Delta lacZ_{\text{Initial}})]$. (C-D) Competitive index scores were calculated as a ratio of output versus input $[(\text{Target}_{\text{Output}}/\Delta lacZ_{\text{Output}}) / (\text{Target}_{\text{Input}}/\Delta lacZ_{\text{Input}})]$. Strains were co-inoculated with a PDH+/PFL+ *ΔlacZ* strain to determine the relative fitness of each test strain. Data for each experiment was obtained from 3 biological replicates.

Supplemental Materials - Methods

*Vibrio cholerae* initial growth on Sigma Type III porcine gastric mucin.
Strains were grown on LB + 0.1mg/mL streptomycin plates overnight at 37ºC and a single colony isolate used to start a fresh LB + 0.1mg/mL streptomycin broth culture grown overnight 210rpm at 37ºC. Overnight cultures were washed in PBS and resuspended to an optical density 1.0 $OD_{600}$. 700µl of either LB, MCLMAN + 0.5% Mucin (Sigma Type III porcine gastric mucin), or MCLMAN minimal media was inoculated 1:1000 with the 1.0 $OD_{600}$ culture. Triplicate 200µl aliquots were dispensed in a 96-well plate and optical densities recorded every 30min for 24h.

Complementation plasmid construction
Complementation plasmid construct open reading frame inserts were generated by PCR using Phusion high-fidelity polymerase (Thermo Scientific). pMMB66EH vector backbones were generated by plasmid purification using Qiagen Mini Prep Kit and subsequent restriction digested using BamHI and HindIII at 37ºC for 1 hour followed by an additional 30 minutes at 37ºC with alkaline phosphatase from calf intestine (CIP) (New England Biolabs). Constructs were assembled using Gibson assembly (New England Biolabs) and were electroporated into electrocompetent S17 λpir *E. coli* and recovered on LB + 0.1mg/mL ampicillin agar plates.

*Vibrio cholerae* complementation strain construction
Complementation strains were made by mating S17 λpir *E. coli* pMMB66EH complementation strains with *ΔaceE*, *ΔaceF* , and *ΔpflA* mutant strains and recovered on LB + 0.1mg/mL ampicillin + 25 U/mL polymyxin B agar plates. Strains were verified using pMMB66EH plasmid-specific primers to detect the full length insert and additional sequencing primers to verify construct sequence.

Complementation growth curves for *ΔaceE* and *ΔaceF*
M9 + 0.5% glucose was prepared and 2mL prewarmed media inoculated 1:250 with 1.0 $OD_{600}$ culture and grown 210rpm at 37ºC. For wild type control and *ΔaceE* complementation strains, 1mM isopropyl-β-D-thiogalactoside (IPTG) was added to induce ectopic expression from the complementation vector. For the *ΔaceF* complementation strain, 0.01mM IPTG was added to induce vector expression. At each timepoint, 100µl was removed for dilution series plating.

Complementation growth curves for *ΔpflA*
M9 + 0.5% glucose 50mM fumarate media was prepared and 700µl was inoculated 1:250 with 1.0 $OD_{600}$ culture. Triplicate 200µl aliquots were dispensed in a 96-well plate and grown statically at 37ºC in anaerobic conditions. Optical densities were recorded every 1h for 24h. For both WT control and pflA complementation strains, 1mM isopropyl-β-D-thiogalactoside (IPTG) was added to induce ectopic expression from the complementation vector.
